# Spatial analysis of human lung cancer reveals organized immune hubs enriched for stem-like CD8 T cells and associated with immunotherapy response

**DOI:** 10.1101/2023.04.04.535379

**Authors:** Jonathan H. Chen, Linda T. Nieman, Maxwell Spurrell, Vjola Jorgji, Peter Richieri, Katherine H. Xu, Roopa Madhu, Milan Parikh, Izabella Zamora, Arnav Mehta, Christopher S. Nabel, Samuel S. Freeman, Joshua D. Pirl, Chenyue Lu, Catherine B. Meador, Jaimie L. Barth, Mustafa Sakhi, Alexander L. Tang, Siranush Sarkizova, Colles Price, Nicolas F. Fernandez, George Emanuel, Jiang He, Katrina Van Raay, Jason W. Reeves, Keren Yizhak, Matan Hofree, Angela Shih, Moshe Sade-Feldman, Genevieve M. Boland, Karin Pelka, Martin Aryee, Ilya Korsunsky, Mari Mino-Kenudson, Justin F. Gainor, Nir Hacohen

## Abstract

The organization of immune cells in human tumors is not well understood. Immunogenic tumors harbor spatially-localized multicellular ‘immunity hubs’ defined by expression of the T cell-attracting chemokines *CXCL10/CXCL11* and abundant T cells. Here, we examined immunity hubs in human pre-immunotherapy lung cancer specimens, and found that they were associated with beneficial responses to PD-1-blockade. Immunity hubs were enriched for many interferon-stimulated genes, T cells in multiple differentiation states, and *CXCL9/10/11*+ macrophages that preferentially interact with CD8 T cells. Critically, we discovered the stem-immunity hub, a subtype of immunity hub strongly associated with favorable PD-1-blockade outcomes, distinct from mature tertiary lymphoid structures, and enriched for stem-like TCF7+PD-1+ CD8 T cells and activated *CCR7*+*LAMP3*+ dendritic cells, as well as chemokines that organize these cells. These results elucidate the spatial organization of the human intratumoral immune response and its relevance to patient immunotherapy outcomes.

## INTRODUCTION

Multicellular networks are critical in mediating immune responses. An important question is how immune cells organize within tumors to effectively eliminate malignant cells. To address this, we previously reported on differences in the baseline immune microenvironment between human mismatch repair deficient (MMRd) and mismatch repair proficient (MMRp) colorectal cancer (CRC). Compared to patients with MMRp, patients with MMRd CRC are dramatically more responsive to immune checkpoint inhibition (ICI)^1^. Based on covarying transcriptional signatures and multispectral imaging of CRC tissues, we observed defined foci of activated T cells expressing *IFNG* abutting malignant and myeloid cells expressing interferon-stimulated genes, including the T-cell-attracting chemokines *CXCL10/CXCL11*^2^. This finding suggests the existence of a positive feedback loop in which activated T cells drive further T cell recruitment.

Cytotoxic CD8 T cells are known to be highly favorable for immunotherapy response across many cancer types^3,4^. Various CD8 T cell differentiation states may contribute to immunotherapy response. Activated PD-1+ and proliferating Ki67+ CD8 T cells are predictive of response^5,6^, suggesting that ongoing CD8 T cell effector activity may be a good predictor of PD-1-blockade responsiveness. Consistent with these findings, prior studies showed that immunotherapy outcomes (including in lung cancer^7^) are positively correlated with IFNγ-related expression signatures^7–9^, including the T cell-attracting chemokine *CXCL10* in melanoma^10^. In addition, the stem-like TCF7+PD-1+ CD8 T cell population has been associated with subsequent patient response to immunotherapy^11–13^, and shown to form a reservoir for the proliferative burst of tumor-specific effector CD8 T cells in mice critical for immunotherapy response^14–16^. However, the spatial organization of these T cell states and their relationships to other cells in the tumor microenvironment are not well understood.

Lung cancer is the leading cause of cancer-related mortality worldwide, and PD-1-blockade is the mainstay ICI therapy, despite only a subset of patients responding to treatment^17^. By performing multiplex RNA- and protein-based imaging, as well as spatial transcriptomics, on tumor tissue from a clinically annotated cohort of immunotherapy-naive non-small cell lung cancer (NSCLC) patients, we here define multicellular networks involved in the anti-tumor immune response – findings that have implications for predicting and augmenting immunotherapy outcomes.

## RESULTS

### Immunity hubs are enriched for T cells and positively associated with response to PD-1-blockade

We previously discovered immunity hubs in MMRd tumors through inference from scRNAseq data and spatial validation using a multiplex panel with RNA *in situ* hybridization (ISH) that consisted of the immunity-hub markers *CXCL10/11* (T cell-attracting chemokines), *IFNG* (T cell activation marker), *CD3E* (T cell marker), *CXCL13* (tumor-reactive T cell marker^18–22^), an antibody cocktail for pan-cytokeratin (PanCK) to mark epithelium, and DAPI (nuclear marker, **Supplementary Fig. 1A-D**).

In the current study, we employed the same panel to test the hypothesis that immunity hubs are associated with response to anti-PD-1 immunotherapy. We collected tissue specimens prior to standard of care anti-PD-1 treatment from a cohort of 68 NSCLC patients, imaged each specimen for the presence of immunity hubs, and tested for association with clinical response. 20 patients demonstrated objective response (complete response [CR] or partial response [PR] by RECIST version 1.1 criteria^23^) and the remaining 48 did not (stable disease [SD] or progressive disease [PD]); additional clinicopathologic characteristics are summarized in **Supplementary Tables 1A and B**.

We first analyzed intratumoral regions, as delimited by the outer border of morphologically neoplastic PanCK+ cells. We defined an immunity hub as the neighborhood around *CXCL10/11*+ cells (by aggregating contiguous 50 x 50μm windows that harbor *CXCL10/11*+ cells; **Fig. 1A-C**), identifying 3777 immunity hubs across 68 samples. We found a 5.5-fold increase of the median frequency of immunity hubs in anti-PD-1 responders relative to non-responders (median of 1.6% of tissue area in responders, 0.30% in non-responders; p=0.0052, **Fig. 1D**). Consistent with our findings in CRC, hubs were enriched in *IFNG*+ (median 4.7-fold increase; median 115 cells/mm^2^ in hubs versus 16 cells/mm^2^ overall; p<1E-6) and *CD3E*+ cells (median 3.0-fold increase; median 782 cells/mm^2^ in hubs versus 204 cells/mm^2^ overall; p<1E-6; **Fig. 1E**).

**Figure 1.**
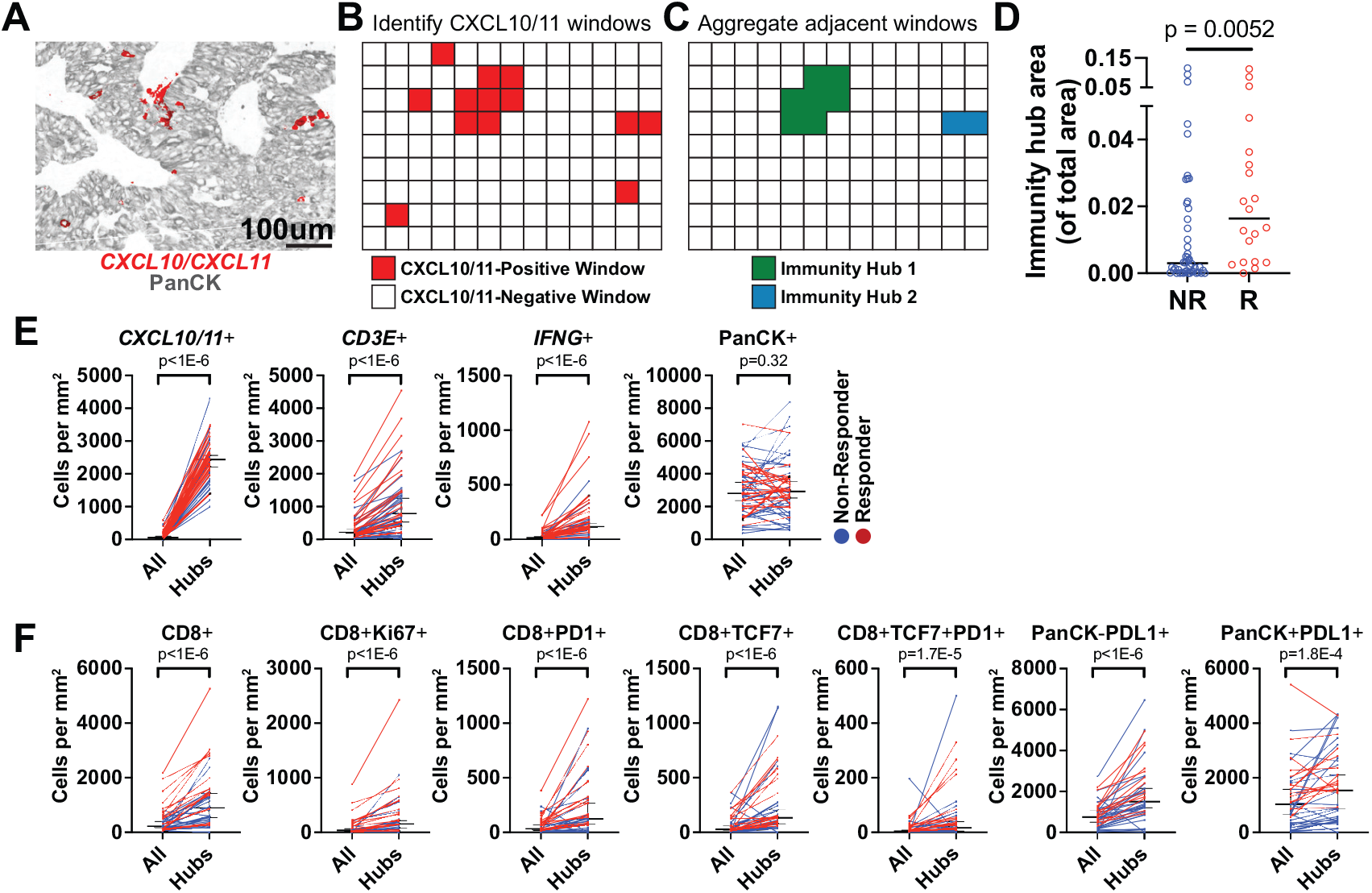
Immunity hubs in tumors and their association with clinical responses to PD-1-blockade in patients with NSCLC. (A) A representative image of a tumor section (Patient #43) stained with the multiplexed RNA ISH/IF panel shows the focal expression pattern of *CXCL10/11* (red). (B) A grid of 50 x 50μm windows was overlaid on images. *CXCL10/11*+ windows (red) were identified using K-means clustering based on *CXCL10/11*+ cell count (where k=2). (C) Adjacent *CXCL10/11*+ windows were aggregated into immunity hubs. Singleton windows were not included as hubs. (D) Immunity hub area as a fraction of total tumor area in responders (PR and CR; n = 20) versus non-responders (SD and PD; n = 48). Mann-Whitney test p-value is shown. (E) Paired comparisons of density of cellular phenotypes from the RNA ISH/IF panel within immunity hubs versus total assessed area, colored by response status (n=59; 1 R and 8 NR tumors lacked immunity hubs). (F) Paired comparisons of density of cellular phenotypes from the IF only panel within immunity hubs versus total assessed area, colored by response status (n=43; 22 samples had images that could not be coregistered and 3 lacked immunity hubs). Each pair of dots connected by a colored line represents a patient. Black horizontal lines denote median and 95% confidence interval. Wilcoxon matched-pairs signed rank test Benjamini-Hochberg (BH) adjusted p-values are shown.

Given that activated CD8 T cells express *CXCR3* (the receptor for CXCL10 and CXCL11)^24^, we hypothesized that activated CD8 T cell populations would be attracted to immunity hubs. We therefore built a multiplexed antibody panel to assess CD8 T cell states (**Supplementary Fig. 1A**) and stained sequential serial tissue sections from the same 68 patient samples (overall enumeration of cellular frequencies by patient in **Supplementary Tables 1C and D**). To integrate our two staining panels, we computationally coregistered 46 paired serial sections (NR=33, R=13) with sufficiently matched regional architecture (**Supplementary Fig. 1E-H**), leading to 2712 observed immunity hubs in the coregistered regions. Immunity hubs harbored a higher density of CD8 T cells relative to the overall tumor (median 2.8-fold increase by paired analysis; median 884 cells/mm^2^ in hubs versus 210 cells/mm^2^ overall; p<1E-6; **Fig. 1F**), including CD8+ T cell substates, such as activated CD8+PD-1+ (median 3.5-fold increase; median 123 cells/mm^2^ in hubs versus 31 cells/mm^2^ overall; p<1E-6) and stem-like CD8+TCF7+PD-1+ cells (median 3.2-fold increase; median 16 cells/mm^2^ in hubs versus 4 cells/mm^2^ overall; p=1.75E-5; **Fig. 1F**). These data demonstrate that immunity hubs are associated with subsequent beneficial responses to PD-1-blockade and are potential foci of CD8 T cell activity.

### The stem-immunity hub represents a subset of immunity hubs enriched for *IFNG* and stem-like CD8+TCF7+PD-1+ T cells and strongly associated with immunotherapy response

We hypothesized that there may be subtypes of immunity hubs with distinct cellular compositions. We performed unsupervised subclustering of the 2712 immunity hubs based on frequencies of cell types, revealing seven subclusters (**Fig. 2A**, each dot on the tSNE plot represents one immunity hub) with representation across multiple patients (**Supplementary Fig. 2A and D**). Subcluster 3 area was markedly elevated in responder patients as a fraction of total area (106-fold, median 1.1% of total area in responders versus 0.011% in non-responders, p=0.00004, **Fig. 2B-D & Supplementary Fig. 2E**). Interestingly, Patient #30 had a high frequency of subcluster 3 (**Fig. 2B**), but was classified as a non-responder by RECIST criteria due to isolated progression in the central nervous system. However, Patient #30 otherwise showed a significant extra-cranial response to therapy that remains ongoing (>5.5 years) despite discontinuing PD-1-blockade after a single-dose due to drug-induced pneumonitis, supporting that subcluster 3 is positively associated with response. Consistently, patients with subcluster 3 representing at least 0.1% of total area experienced superior progression-free survival (PFS, p=0.0035) and overall survival (OS, p=0.017) relative to patients below this threshold (**Fig. 2E-F** and **Supplementary Fig. 2F-G**). Compared with the other subclusters, subcluster 3 was enriched for *IFNG*+ cells (median 2.4-fold increase in paired analysis, median 280 cells/mm2 in subcluster 3 versus 91 cells/mm2 in other subclusters, p<0.0001), non-epithelial PanCK-TCF7+ cells (median 4.7-fold in paired analysis, median 1369 cells/mm2 in subcluster 3 versus 278 cells/mm2 in other subclusters, p<0.0001), and most notably, stem-like CD8+TCF7+PD-1+ cells (median 9.7-fold in paired analysis, median 114 cells/mm^2^ in subcluster 3 versus 7 cells/mm^2^ in other subclusters, p<0.0001, **Fig. 2G**, **Supplementary Fig. 2H and I**). The density of CD8+TCF7+PD-1+ cells outside of hubs in these tumors was just 6 cells/mm^2^, revealing subcluster 3 immunity hubs as distinct niches for stem-like CD8 T cells within the entire tumor microenvironment. Inspection of images in locations assigned to subcluster 3 revealed dense aggregates of TCF7+ cells, including CD8+TCF7+PD-1+ cells (**Fig. 2H and Supplementary Fig. 3**). Given the high frequency of stem-like CD8+TCF7+PD-1+ cells together with *IFNG*+ cells, we henceforth refer to subcluster 3 as a “stem-immunity hub.”

**Figure 2.**
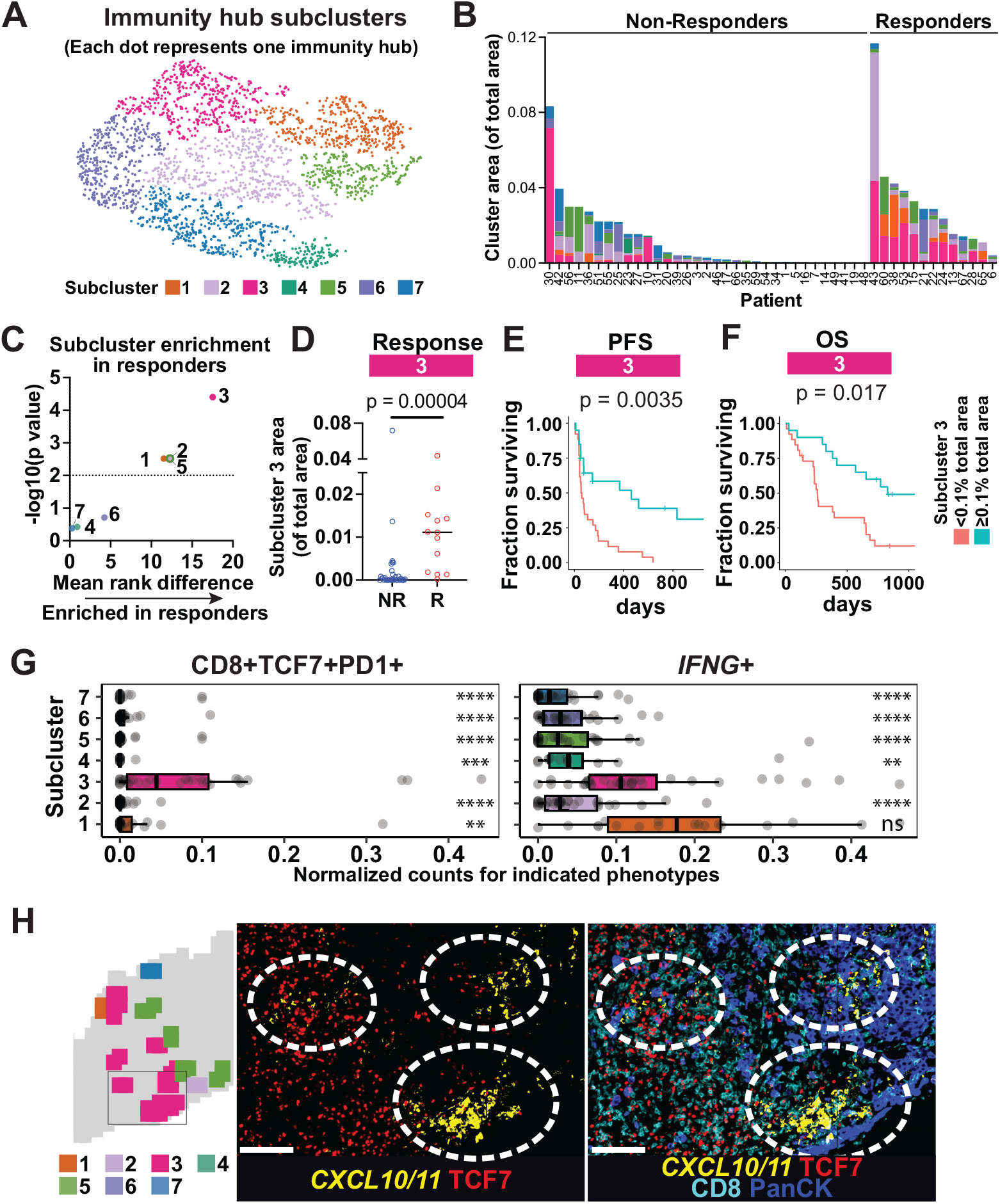
A subcluster of immunity hubs with *IFNG*+ cells and CD8+TCF7+PD-1+ T cells and association with response to PD-1-blockade. (A) Leiden analysis of immunity hubs reveals 7 subclusters. Each point on the tSNE plot represents one immunity hub. (B) Leiden subcluster composition by patient, plotted by the proportion of tumor area in each subcluster. (C) Mean rank difference and associated BH adjusted p-values calculated for each Leiden subcluster by comparing responders versus non-responders. Values are derived from the comparisons shown in Supplementary Fig. 2E. (D) Immunity hub subcluster 3 is enriched in responders. The proportion of tumor area that is in immunity hub subcluster 3 is shown for each patient. BH adjusted Mann-Whitney test p-value is shown. (E-F) Immunity hub subcluster 3 is associated with increased progression-free survival (PFS; E) and overall survival (OS; F) by Kaplan-Meier analysis. Patients were classified as either above or below a threshold of 0.1% of total tumor area for subcluster 3. Units of time are in days. BH adjusted p-values are shown. (G) Immunity hub composition by subcluster for the indicated phenotypes. Phenotype counts were normalized by immunity hub area and scaled from 0-1. Each point represents the mean value across all immunity hubs of a given Leiden subcluster for each patient having that subcluster. Statistical comparison was performed using an unpaired Mann-Whitney test relative to subcluster 3. Benjamini-Hochberg adjusted p-values shown: ns = not significant, *p<0.05, **p<0.01, ***p<0.001, and ****p<0.0001. (H) Spatial map of tissue from a responder (Patient #28) showing immunity hubs color-coded by Leiden subclusters (left). A region containing three subcluster 3 immunity hubs is indicated by a box in the spatial map. Images (center and right) from the same boxed inset region are shown with the indicated markers. White scale bar denotes 100um.

### Stem-immunity hubs overlap with aggregates of PanCK-TCF7+ cells, express CCR7 ligands and are distinct from mature TLS

Given the striking abundance of TCF7+ cells in immunity hub subcluster 3, we further tested whether the observed strong association with response was attributable to aggregates of TCF7+ cells, irrespective of *CXCL10/11* expression. To define aggregates of TCF7+ cells, we again employed a 50 x 50μm grid on coregistered images, using PanCK-TCF7+ cellular densities as input for K-means clustering (k=2, **Fig. 3A**). We observed that the frequency of TCF7+ aggregates (**Fig. 3B**) only weakly trended with response. However, stem-immunity hub subcluster 3 (which strongly associated with response) was enriched for proximity to TCF7+ aggregates, relative to the other subclusters (**Fig. 3C-D** and **Supplementary Fig. 4A**). Thus, immunity hubs overlapping with TCF7+ aggregates are overwhelmingly those in subcluster 3.

**Figure 3.**
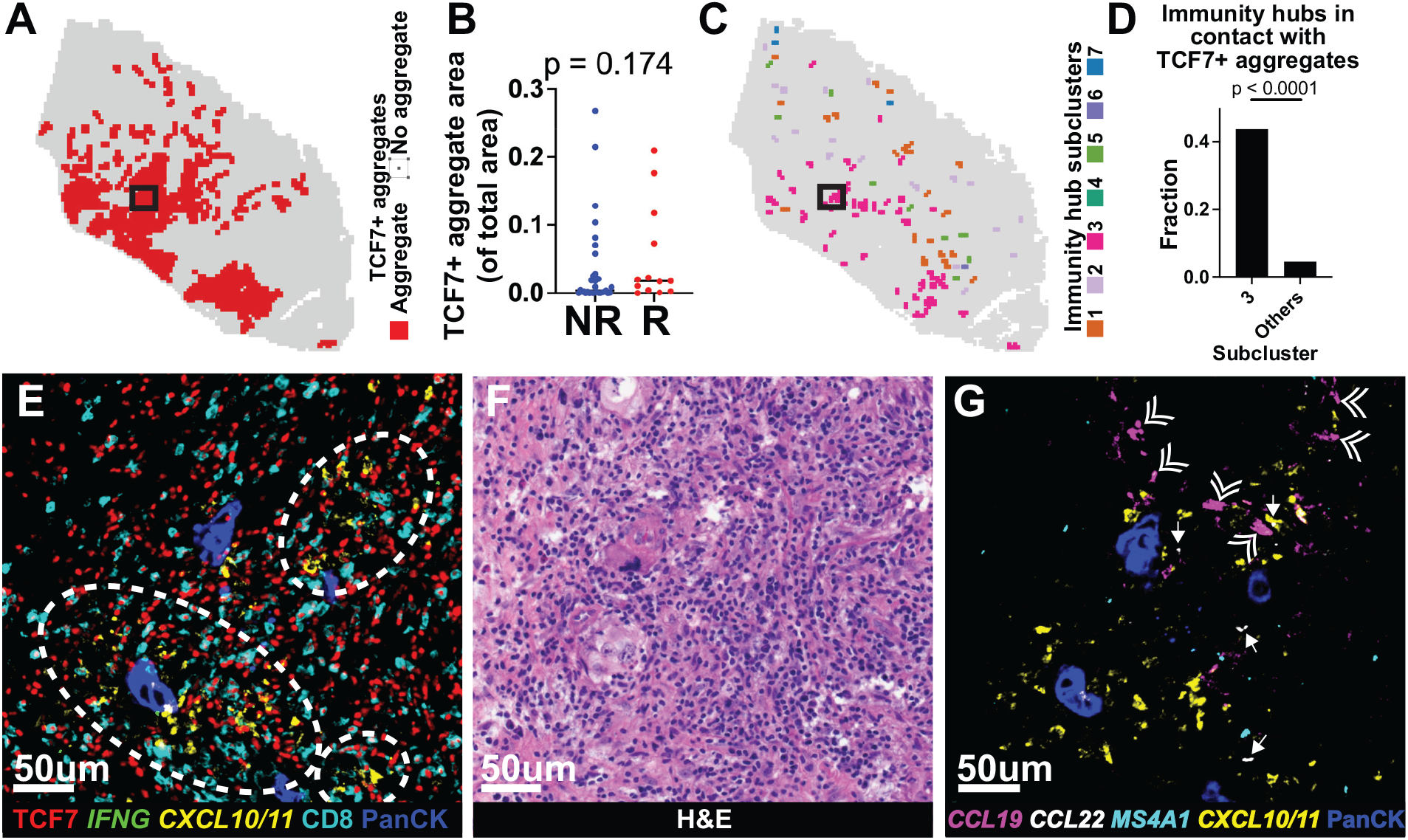
Overlap of immunity hub subcluster 3 with TCF7+ aggregates. (A) Spatial map (from Patient #53) showing TCF7+ aggregates identified by K-means clustering with k=2 based on PanCK-TCF7+ cell count. (B) TCF7+ aggregate area (as a fraction of all area) plotted by response status. Statistical comparisons were performed using an unpaired Mann-Whitney test. (C) Spatial map of tissue from patient in (A) showing immunity hubs (from Fig. 2) color-coded by Leiden subcluster. (D) Frequency of contact (at least touching in a cardinal direction) between TCF7+ aggregates and subcluster 3 immunity hubs versus all other immunity hub subclusters (i.e. subclusters 1-2 and 4-7). Statistical comparison was performed using Fisher’s exact test on counts shown in Supplementary Fig. 4A. (E) Coregistered image of immunity hub subcluster 3 (from region in Patient #53 denoted by box in (A and C)). Dashed ovals indicate areas with *CXCL10/11*+ cells. (F) Hematoxylin and eosin (H&E) stained serial section from the area matched to that in (E). (G) Multiplexed RNA ISH/IF image of serial section from area matched to that in (E). Double arrowheads indicate *CCL19*+ cells. Arrows indicate *CCL22*+ cells.

Since tertiary lymphoid structures (TLS) harbor TCF7+ cells^25^, we tested whether stem-immunity hubs were TLS. TLS are ectopic lymphoid aggregates that develop in chronically inflamed tissue and mature to form B cell follicles and germinal centers^26^ visible by light microscopy. To examine the presence of B cells, we stained sequential sections from six cases (NR=2, R=4) using a third panel (*MS4A1* for B cells along with *CXCL10/11*, and PanCK). TCF7+ aggregates that contained *CXCL10/11+* cells (i.e. stem-immunity hubs) lacked B cell follicles (**Fig. 3E-G and Supplementary Fig. 4B-D**). These results suggest that stem-immunity hubs are compositionally distinct from mature TLS (because they lack B cell follicles).

We reasoned that because stem-like TCF7+PD-1+ CD8 T cells have been reported to express CCR7^27–29^ and migrate in response to the CCR7 ligands CCL19/21^28^, stem-immunity hubs might express CCL19, despite the absence of B cell follicles. Indeed, examined regions in stem-immunity hubs showed *CCL19* expression, which in some cells was co-expressed with *CCL22* (**Fig. 3G** and **Supplementary Fig. 4D**, responder cases), previously shown to be a marker for “mreg” dendritic cells^30^. Since aggregates featuring TCF7+ and *CXCL10/11*+ cells also express *CCL19*, we conclude that stem-immunity hubs feature a confluence of two major chemokine pathways (CXCR3 and CCR7 ligands), but are not mature TLSs.

### Stem-immunity hubs are enriched in IFNγ-induced transcripts, macrophages, and activated T cells and depleted in B cells relative to TLS

To quantitatively examine the differences between stem-immunity hubs and TLS, we profiled ~1800 genes using spatially-indexed transcriptomic analysis (GeoMx^31^) in five immunotherapy-naive primary human NSCLC tumor samples. Similar to laser-capture microdissection, GeoMx enables gene expression readouts of user-defined spatial regions of interest. Within each GeoMx slide, we used a combination of a nuclear stain to find nuclear aggregates and *CXCL10* and *CCL19* RNA FISH probes to define stem-immunity hubs (*CXCL10+CCL19*+), non-stem-immunity hubs (*CXCL10+CCL19*−), and TLS-like structures (*CXCL10-CCL19*+) (see **Supplementary Fig. 5A** for criteria; average ROI area = 56,326um^2^; 238 total ROI profiled). Each slide featured many instances of all three structures.

We identified genes that were differentially expressed between stem-immunity hubs and TLSs (**Fig. 4**), and found that stem-immunity hubs were strongly enriched for the “Interferon Gamma Response” gene set^32^ relative to TLS (adjusted p-value = 3.42×10^−7^; **Supplementary Table 1E**), including *CXCL10* (used to select ROIs) and its close homolog *CXCL9*. Stem-immunity hubs were also enriched for markers consistent with activated CD8 T cells (e.g. *CCL5*, *CD8A*, *CD2*, and *LAG3*) and macrophages (e.g. *C1QA*, *C1QB*, *LYZ*, *CD14*, *CD68*, *CD163*, *S100A9*, *CCL18*, *CSF1R*, and *TLR2*^2^). In contrast, TLS were significantly enriched for B cell genes (e.g. *CD19*, *MS4A1*, *CR2*, *CD22*, *TNFRSF13C*, *BLK*, *CD79A*, and *CD79B*^2^) and modestly enriched for markers associated with less-differentiated lymphocyte states (*TCF7, CCR7*, and *SELL*^2,14,33^).

**Figure 4.**
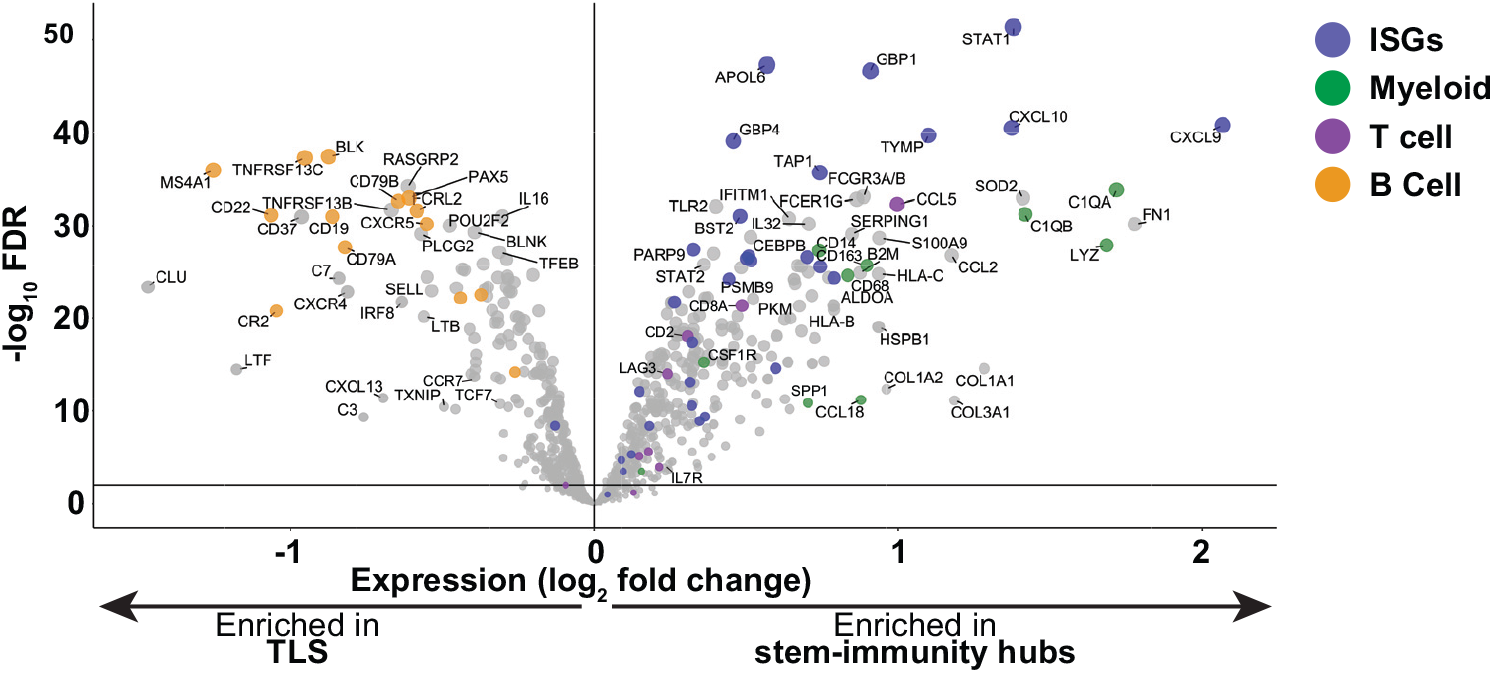
Expression of T cell and macrophage transcripts in stem-immunity hubs and TLS. Comparison of stem-immunity hub and TLS (as defined in Supplementary Fig. 5A) gene expression using GeoMx CTA assay. Volcano plot depicts pooled ROIs across all 5 samples. Each dot represents one gene. Dot coloring is as follows: blue - ISGs, green - myeloid genes, purple - T cell genes, and orange - B cell genes.

As expected from our ROI selection strategy (**Supplementary Fig. 5A**), stem-immunity hubs expressed substantially more *CCL19* than non-stem-immunity hubs (**Supplementary Fig. 5B** and **Supplementary Table 1F**). Consistent with our observation that stem-immunity hubs contained lymphoid aggregates, stem-immunity hubs were enriched in T and B cell markers associated with less-differentiated lymphocytes (*TCF7*, *SELL*, *IL7R*, and *CCR7*, **Supplementary Fig. 5B**). In contrast, non-stem-immunity hubs were modestly enriched for a subset of interferon stimulated genes (ISGs) and macrophage markers (**Supplementary Fig. 5B**). We conclude that the stem-immunity hub is distinct from the B cell-rich mature TLS based on elevated levels of interferon-stimulated genes and genes marking macrophages and T cells.

### Spatial transcriptomics at single-cell resolution reveals gene expression, cellular composition and cell-cell interactions within stem-immunity hubs

As an independent approach to identify spatially organized hubs within NSCLC tumors, multiplexed error-robust FISH (MERFISH^34^) was used for single-cell resolution detection of 479 genes that mark immune, stromal, and epithelial cell states (**Fig. 5A**). We selected four NSCLC cases with *CXCL10/11*+ expression and large tissue area from patients who subsequently responded to PD-1-blockade. We divided the tissue into 2500um^2^ tiles (69,097 total tiles; median 2553 transcripts and 288 genes detected per tile) and applied Louvain clustering based on differential gene expression, revealing 14 tile clusters (**Supplementary Fig. 6A-D**), including: stem-immunity hubs (elevated *CCL19* and *CXCL9/10/11;* cluster t7; **Supplementary Fig. 6D and F**), non-stem-immunity hubs (elevated *CXCL9/10/11* but not *CCL19;* cluster t4), vascular hubs (expressing endothelial markers *VWF* and *ERG* and pericyte markers *CSPG4* and *GJA4*; clusters t2 and t8), and all other regions, broadly labeled “tumor” (t0, 1, 3, 5, 6, 9-13; see **Supplementary Table 1H** for comparison of four types of tile clusters). The definition of stem-immunity and non-stem-immunity hubs in our MERFISH and GeoMx dataset are in agreement based on concordant expression of 177 genes measured by both technologies (r=0.78, p<1E-35; **Supplementary Fig. 6G**).

**Figure 5.**
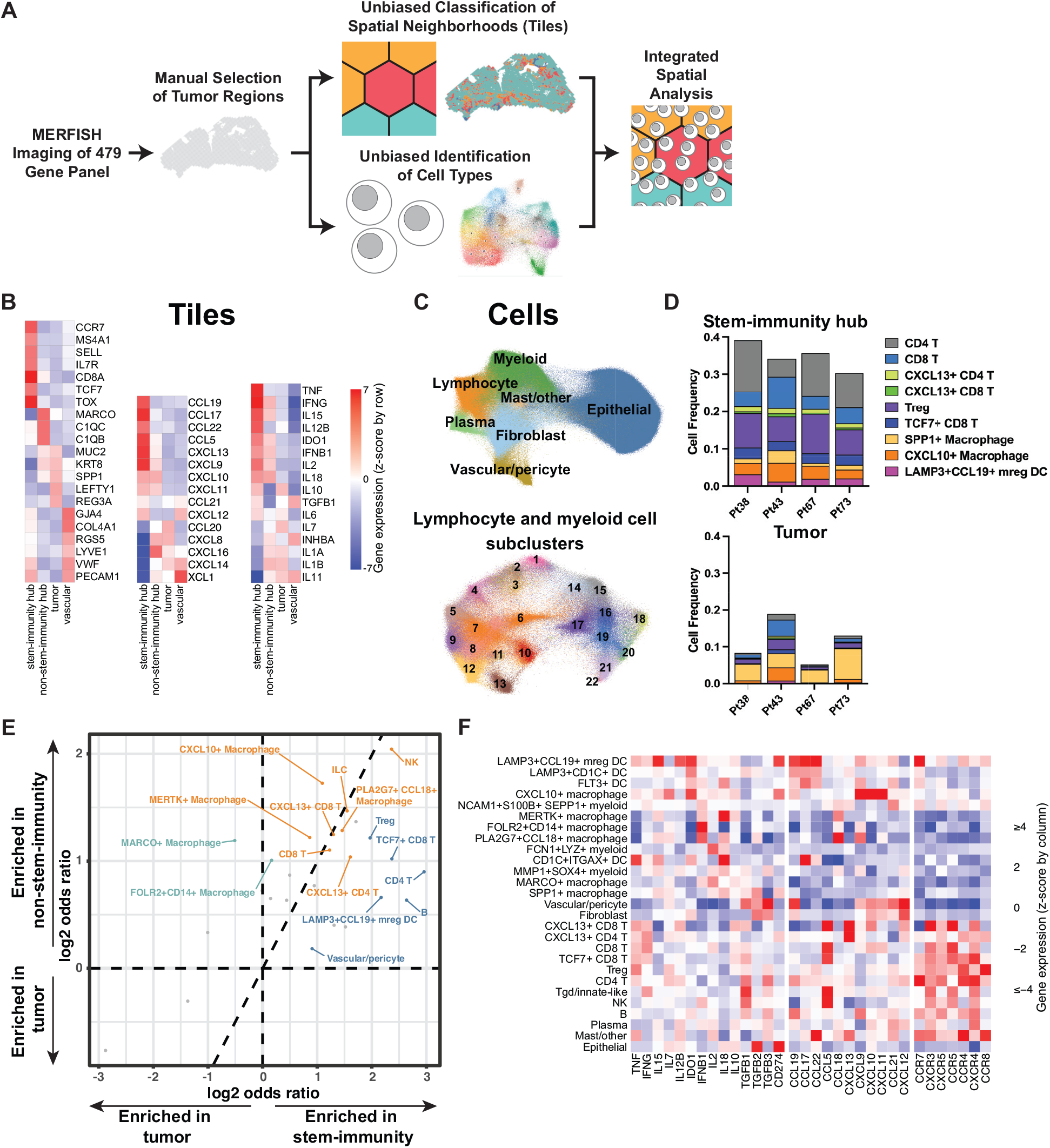
MERFISH spatial analysis of lymphocyte and myeloid populations in immunity hubs. (A) Schematic of MERFISH analysis. Tissue was manually annotated to include tumor areas, but exclude non-neoplastic areas, necrotic regions, and tissue folds. Two non-mutually-exclusive approaches were taken for analysis, assigning transcripts to: (top) tiles within a regular grid and (bottom) segmented cells. Clustering analysis was performed for both tiles and cells. Furthermore, every cell could be uniquely assigned to one tile. (B) Expression of indicated genes for marker genes (left), chemokines (center), and other secreted factors (right) in indicated tile clusters. Tile identities were assigned as shown in Supplementary Fig. 6. (C) UMAP of coarse cell type clusters (top) and finely clustered immune cells (bottom) from MERFISH data. Each dot represents one cell. For finely clustered immune cells, clusters are as follows: 1. *LAMP3+CCL19*+ mreg DC, 2. *LAMP3+CD1C*+ DC, 3. *FLT3*+ DC, 4. *CD1C+ITGAX*+ DC, 5. *FCN1+LYZ*+ myeloid, 6. *CXCL10*+ macrophage, 7. *MARCO*+ macrophage, 8. *FOLR2+CD14*+ macrophage, 9. *MERTK*+ macrophage, 10. *MMP1+SOX4*+ myeloid, 11. *PLA2G7+CCL18*+ macrophage, 12. *SPP1*+ macrophage, 13. *NCAM1+S100B+SEPP1*+ myeloid, 14. B cells, 15. CD4 T cells, 16. *TCF7*+ CD8 T cells, 17. Treg, 18. *CXCL13*+ CD4 T cells, 19. CD8 T cells, 20. *CXCL13+* CD8 T cells, 21. Tgd/innate-like, and 22. NK. (D) Frequency of indicated cell types within stem-immunity (upper) and tumor (lower) tile neighborhoods. Each column represents the indicated patient. (E) Log2 odds ratio (OR) enrichment of the abundance of indicated cell populations within hubs: stem-immunity versus tumor (x-axis) or non-stem-immunity versus tumor (y-axis). Populations are called out as significantly enriched (FDR<20%, logOR>0) in non-stem-immunity hub (teal), stem-immunity hub (blue) or both hub types (orange). (F) Expression of indicated genes by cell type, subsetted on cells located within stem-immunity hubs.

We sought to determine which chemokines and cytokines are present in stem-immunity hubs that might attract and support stem-like T cells (**Supplementary Table 1H**). Stem-immunity hubs were indeed enriched for T cell-attracting chemokines relative to tumor, including *CCL19*, *CXCL9*, *CXCL10*, *CXCL11, CCL17*, *CCL22*, and *CCL5* (**Fig. 5B**). We noted increased expression of *IL15* and *IL12B* in the stem-immunity versus tumor hubs, both of which may support T cell survival and differentiation (**Fig. 5B**). Consistent with higher density of activated immune cells, there was also increased expression of *IFNG*, *TNF*, *IL2*, and *IDO1* in stem-immunity hubs relative to tumor.

We then examined the cell type composition of stem-immunity and non-stem-immunity hubs to identify the relevant multicellular networks involved in hub activity. We identified and labeled cells with our internal multi-sample MERFISH analysis pipeline, which combined cell segmentation, quality control, graph-based clustering, and mixed effects differential expression. With this approach, we identified 1,488,870 cells across four samples (**Fig. 5C**), with median 36 transcripts and 24 genes per cell, annotated into 7 lineages (**Supplementary Fig. 7A-C and Supplementary Table 1I**) distributed among the four major tile clusters described above (**Supplementary Table 1J**). To better resolve the immune cell composition, we performed fine-grained clustering of lymphocytes and myeloid cells (**Fig. 5C; Supplementary Fig. 7D-F and Supplementary Table 1K**).

Stem-immunity hubs (tile cluster t7) were enriched for *TCF7*+ CD8 T cells, as expected (**Fig. 5D-E**), as well as CD8 and CD4 T cells (**Fig. 5D-E**), but depleted for tumor cells (**Supplementary Fig. 8A and B** and **Supplementary Table 1J**), consistent with our antibody staining (**Supplementary Fig. 2H and I and 3** and **Figs. 2H and 3E**). Among myeloid cells, there was enrichment for activated “mreg” DCs, which present antigen to T cells (**Fig. 5D-E and Supplementary Table 1J**), and depletion for *SPP1+* macrophages (**Supplementary Fig. 8A and B**), which have been associated with fibrosis and poor patient outcomes^35^. Focusing on cytokines and chemokines, we found that the mreg DC population expressed *CCL19*, *CCL17*, *CCL22*, *IL15*, *IL12B*, and *IDO1* (**Fig. 5F** and **Supplementary Table 1K**), consistent with published single-cell signatures^2,30,36^. These findings suggest that the stem-immunity hub is relatively spatially separated from tumor cells and well-equipped to support T cells through molecules such as IL15^37,38^.

To better understand cell-cell interactions in the stem-immunity hub, we asked which cells are colocalized with *CCL19+* mreg DCs and *CXCL10*+ macrophages, as these cells produce the chemokines which define the stem-immunity hub. We found that mreg DCs were most frequently adjacent to conventional CD4 T cells (FDR = 1×10^−5^) and Tregs (FDR = 5.7×10^−6^; **Fig. 6A and B, Supplementary Fig. 9**, and **Supplementary Table 1M**), consistent with *CCR7* (receptor for CCL19) and *CCR4* (receptor for CCL17/22^39^) expression on CD4 T cells (**Fig. 5F** and **Supplementary Table 1K**) and with independent reports that mreg DCs preferentially interact with CD4 T cells (including Tregs)^40,41^. These data suggest that stem-immunity hubs might be sites for activation of CD4 T cells as cytotoxic effectors^42^, or helper T cells^43^ or Tregs^41^ that can license or inhibit DCs to activate CD8 T cells, respectively. *CXCL10*+ macrophages preferentially interacted with CD8 T cells within and outside stem-immunity hubs (**Fig. 6A and B** and **Supplementary Fig. 9**), in agreement with a recent study finding this pair of cells forming stable doublets by flow cytometry^40^. These macrophages may promote CD8 T cell exhaustion, as was recently shown^44,45^. Thus, the stem-immunity hub contains two axes of myeloid-T cell interactions, mreg DCs with CD4 T cells and *CXCL10+* macrophage with CD8 T cells (**Fig. 6C**).

**Figure 6.**
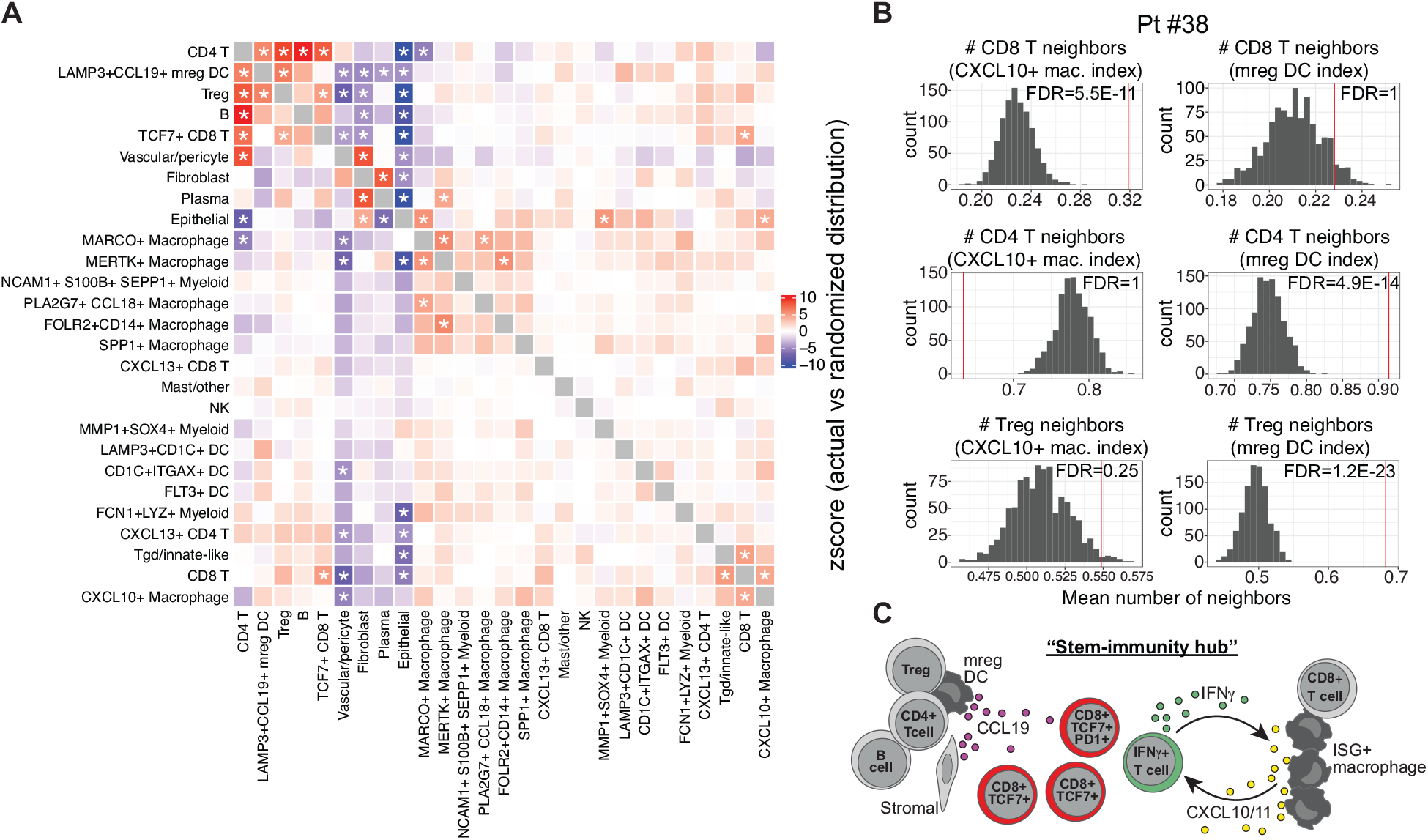
Cell-cell interactions within stem-immunity hubs. (A) Neighboring cell analysis for cells within stem-immunity hubs. For each plot, indicated cluster position (row) is held constant and enrichment with partner (column) is quantified versus scrambled distribution. Mean z-scores across all four samples are shown. Two-tailed p-values were calculated from the mean z-scores and FDRs computed with the Benjamini-Hochberg procedure across all p-values. FDR<0.05 are denoted with an asterisk. (B) Red line on histograms show how frequently specific T cell types are observed to neighbor the indicated myeloid cell type within the stem-immunity hubs for Patient #38. Gray bars denote random distribution (expected distribution based on cell type frequency). (C) Graphical schematic of the stem-immunity hub.

## DISCUSSION

Our study characterizes spatially-organized immunity hubs and identifies a new subset of these hubs, the stem-immunity hub, in human lung tumors. Distinct from mature TLS, the stem-immunity hub is defined by presence of stem-like *TCF7+* CD8 T lymphocytes and the expression of the chemokines *CXCL10/11* and *CCL19*, with CD8 T cells adjacent to *CXCL10+* macrophages and CD4 T cells adjacent to *CCL19+* mreg DCs. The presence of immunity hubs, and especially stem-immunity hubs, in pre-therapy biopsies was associated with subsequent beneficial responses to anti-PD-1 immunotherapy in patients with advanced lung cancer.

Immunity hubs likely represent the major site for aggregation of cells expressing interferon-stimulated genes^8,9,46–48^, including CXCR3 ligands CXCL9/10/11, in melanoma^7,10^, CRC^49^, breast, ovarian and other tumor types^7^. These genes are critical for effective tumor immunity and were shown in numerous studies to be associated with PD-1-blockade responses in patients^2,50–52^.

The stem-immunity hub might contextualize recent reports regarding stem-like T cells within the tumor microenvironment. First, stem-like CD8 T cells were found in aggregates^11^ and near blood vessels, but away from tumor cells^14,38^. Second, signals from DCs within the tumor microenvironment have been shown sufficient to differentiate stem-like T cells (newly arrived from lymph nodes) into effector T cells^53,54^.

The organization of the stem-immunity hub is reminiscent of the lymph node T cell regions (paracortex and interfollicular zone) with respect to cellular composition. In particular, the lymph node T cell regions not only houses TCF7+ T cells, but also uses CCL19 to position DCs and CXCL9/10 to position T cells^55,56^. Since mreg DCs within stem-immunity hubs are likely to be more enriched for tumor antigens compared to migrating DCs present in lymph nodes, the local tumor hubs could be superior to lymph nodes in expanding tumor-reactive PD-1+TCF7+ stem-like T cells. Conversely, the lymph node may specialize in preserving the most primordial tumor-reactive stem-like T cells^54^ before their exit to the periphery and entry into the tumor.

In conclusion, our data support a central role for *CXCL10/11*-expressing immunity hubs in anti-tumor immunity and responsiveness to PD-1-blockade in NSCLC patients. Furthermore, stem-immunity hubs represent a subclass of immunity hubs enriched for TCF7+PD-1+ CD8 T cells and *CCL19,* and are particularly associated with clinically beneficial responses. How the hubs change after treatment with checkpoint blockade remains unknown given the challenge of attaining research biopsies from responding NSCLC patients. These multicellular networks in human lung cancer are promising candidates for predictive assays and represent potential therapeutic targets for cancer immunotherapy.

## ONLINE METHODS

### Patient cohort

Formalin-fixed paraffin-embedded (FFPE) whole-tissue sections from 68 patients with NSCLC treated with PD-1/PD-L1 inhibitor therapy were retrospectively collected from Massachusetts General Hospital. The cohort was 47% female, with an average age of 69 years-old, and 46% first-line therapy; additional baseline clinical and pathologic characteristics of the cohort are summarized in **Supplementary Tables 1A** and **B**. Response to therapy was assessed using RECIST criteria (version 1.1^23^). Progression-free survival was measured from the date of PD-1-blockade initiation to radiographic disease progression or death. Overall survival was measured from the date of therapy initiation to the date of death. Patients still alive at the date of cutoff were censored (final data cutoff February 7, 2022)(**Supplementary Table 1B**). Clinical IHC for PD-L1 (Cell Signaling Technologies clone E1L3N) was performed by the MGH Department of Pathology and evaluated as the percentage of tumor cells with any degree of PD-L1 expression, as previously described^57^. Pre-treatment FFPE tissue specimens were retrieved from the MGH Department of Pathology clinical archive and evaluated for presence of tumor by a board-certified pathologist (JHC) and a minimal evaluable tumor area of 0.5 mm^2^. 5um sections were cut by the MGH Histopathology Research Core facility onto standard positively charged slides. This study was conducted under MGH IRB protocol #2019P002829 and DF/HCC 02-240.

### Tissue staining

Tissue sections were stained with Leica Bond RX automated stainer using unlabeled primary antibodies and tyramide signal amplification-based Opal fluorophores (Akoya Biosciences). The RNA ISH/IF panel also employed RNAscope ISH probes (Advanced Cell Diagnostics). Reagents are described in **Supplementary Table 1N**. The construction of the RNA ISH/IF panel has been described^2^. The IF only panel was constructed in accordance with the Akoya Phenoptics Opal Assay Development Guide. Primary antibodies were titrated in tonsil and lung cancer tissue. Multiplex staining employed the Opal 7-Color Automation IHC Kit (NEL821001KT, Akoya Biosciences). For PD-1 and PD-L1 staining, Powervision Poly-HRP anti-Rabbit IgG (PV6119, Leica Biosystems) at 1:10 dilution was employed because of superior signal to background performance. Fluorophore concentrations were selected based on balancing signal intensity across neighboring channels. The absence of bleedthrough was confirmed by fluorescence minus one experiments. Importantly, fluorophores were selected for maximum compatibility with the MOTIF setting on the Vectra Polaris, which allows for the use of fluorophores that are widely spectrally separated. Once the staining panel was locked, samples were run in batches of 10-18 slides. Each run contained a mixture of responder and non-responder samples. A tonsil slide was included in all batches as a control. For the RNA ISH/IF panel, a DNA mismatch repair deficient colon cancer slide was also included as a control^2^.

### Image acquisition

Whole slide images were acquired on an Akoya Vectra Polaris multispectral slide scanner at 20x (0.5μm) resolution. Spectral unmixing was performed using inForm (Akoya Biosciences). The inForm unmixing library was constructed using single color control tonsil slides (CD20 or DAPI) and an unstained lung cancer slide for autofluorescence control. Cell segmentation and quantification was performed using Indica Labs Halo software on tumor regions identified by a board-certified pathologist (JHC). Areas within tumor regions containing low quality regions such as folds, necrosis, or staining artifacts were excluded from the analysis.

### Cell Phenotype Enumeration

Image analysis was performed blinded to response status. Cell segmentation and phenotyping of whole slide images from 68 patients was performed using the FISH-IF v.1.2.2 Halo module for the RNA ISH/IF panel and using the HighPlex FL v3.2.1 Halo module for the IF only panel. Nucleus-based cell segmentation was performed using DAPI. The cell boundary was delineated by dilating nuclear objects 2-3μm.

For the IF only panel, cells were phenotyped based on intensity thresholds and percentage coverage of stain within the nuclear and cytoplasmic compartments. Percentage coverage was utilized to minimize effects of intensity overlap from bright neighbors. Cells were considered positive for a particular phenotype (e.g. CD8) if there was positivity in either compartment. For the nuclear factors, TCF7 and Ki67, only the nuclear intensity was used to determine positivity.

For the RNA ISH/IF panel, cell positivity for ISH markers was set using criteria for stain intensity and size, where brighter and larger RNA dots were given more weight. All markers were required to be negative for autofluorescence. For both panels, analyzed samples were manually checked for quality of phenotyping. To assess for false positive calls, 100 cells called positive for a given phenotype were manually reviewed to confirm they were true positives. Particular emphasis was placed on the cells that were just above the minimum threshold for a positive call (although we chose not to analyze our positive cells by intensity substratification beyond this QC, the analysis software outputs intensity values by default). To assess for false negative calls, 100 cells called negative were similarly checked. Additionally, cells visually appearing positive were checked for consistency of calls. This approach was performed for all markers for each case. Object tables containing individual cell information from all cell phenotyping analyses were exported for further analysis using the R programming language.

### Combined panels

Matched serial sections from 46 patients stained from the two panels (RNA ISH/IF and IF only) were coregistered using Halo (Indica Labs) software. The 22 samples that could not be coregistered were generally small and/or had regional architecture substantially different between the two slides, which made finding a registerable region of tumor impossible. Landmarks were identified in each image to ensure good alignment of tumor regions. Merged images underwent cell segmentation using the machine learning default AI module in Halo software. Cell phenotyping was performed using the FISH-IF v.2.1.5 Halo module for the RNA ISH/IF panel and using the HighPlex FL v4.1.3 Halo module for the IF only panel. All object tables were exported for analysis in R.

### Unbiased window identification

A computationally fast window approach was used to explore the local tissue microenvironment, where cell phenotypes were enumerated within a 50 x 50μm window. Then, an unbiased K-means clustering was applied across all windows using the *CXCL10/11*+ cell fraction (of all cells within a window). Windows that were enriched in *CXCL10/11*+ cells grouped into cluster 2 (i.e. immunity hub windows), while windows that had little to no *CXCL10/11+* cells were assigned to cluster 1. An analogous approach was taken for PanCK-TCF7+ cells. Immunity hub windows were overlaid onto a map of the tissue allowing visual inspection of these regions.

### Window combination and tSNE analysis

Our windowing approach divided large areas enriched in *CXCL10/11*+ cells into multiple immunity hub windows. Therefore, adjacent *CXCL10/11+* windows were combined to facilitate a compositional analysis more reflective of the integrative biology across the entirety of the immunity hub. This allows us to capture a community of interacting cells that span multiple grid windows. An immunity hub is defined as two or more *CXCL10/11+* windows that are adjacent and each had greater than 7 cells (15 cells for the two coregistered images). For the 68 coregistered images, we had the following total counts: 1,330,963 total windows (of which 1,144,589 had >7 cells), 22,028 *CXCL10/11*+ windows (of which 20,387 had >7 cells), and 14,139 CXCL10/11+ windows in the 3,777 aggregates. For the 46 coregistered images, we had the following total counts: 865,900 total windows (of which 688,172 had >15 cells), 15,963 CXCL10/11+ windows (of which 14,083 had >15 cells), and 9,939 *CXCL10/11*+ windows in the 2712 aggregates. Each hub was assigned a unique identifier. For each immunity hub, cell phenotypes were enumerated and the area was computed. For each patient sample, immunity hubs were mapped back onto a plot of the tissue section for visual inspection. A similar aggregation approach was taken for defining TCF7+ aggregates. An expanded search window (100um) was employed to fully agglomerate large contiguous aggregates. Each aggregate was required to be composed of two or more PanCK-TCF7+ neighboring windows and have a minimum of 35 cells.

Paired plots were generated comparing phenotype density in immunity hubs compared to all tumor area (both immunity hub and non-immunity hub). Benjamini-Hochberg adjusted p-values for the paired Wilcoxon test were calculated (4 phenotypes compared in RNA panel and 7 in antibody panel).

In order to explore cell phenotype composition within the 2712 immunity hubs, we performed unsupervised clustering analysis on the hubs using the Rtsne package (approach adapted from Ref^58^). Cell phenotype counts per hub area were scaled based on the global maximum for each phenotype and used to create tSNE coordinates. 21 cell phenotype features were used in the analysis: *CD3E+,* CD8+, CD8+Ki67+, CD8+PD-1+, CD8+PD-1+Ki67+, CD8+PD-1+TCF7+, CD8+TCF7+, CD8+TCF7+Ki67+, PanCK+ (IF only panel), PanCK+PD-L1+, PanCK+TCF7+, PanCK-PD-L1+, PanCK-TCF7+, PanCK+ (RNA ISH/IF panel), PanCK+*CXCL10/11*+, PanCK-*CXCL10/11*+, *CXCL10/11+*, *IFNG+*, PD-1+, PD-L1+, and total cell density. The optimal number of principal components for variance explained was identified as 6 by scree plot. Thereafter a tSNE (t-distributed Stochastic Neighbor Embedding) dimensionality reduction was performed using Rtsne, with perplexity set to 50. To identify subclusters, the tSNE coordinates were used to create an edge list for a fast k-nearest neighbor search algorithm with the maximum number of nearest neighbors equal to 50. Next, distances were converted to weights such that greater weight was given to smaller neighbor distances. A Leiden clustering algorithm^59^ was then applied using the modularity method with the resolution parameter set to 0.25, yielding 7 Leiden subclusters. The subclusters were mapped back onto a tissue plot for closer inspection of cell phenotypes in these regions. Plots were also generated showing subclusters present within each patient image and the cell phenotype composition per subcluster. P-values were adjusted for multiple hypothesis testing using the Benjamini-Hochberg method.

Computation was performed in R using the packages tidyverse, data.table, Rtsne, FNN, igraph, stats, ggpubr, Rstatix, factoextra, gridExtra, ggforce, RColorBrewer, survminer.

### GeoMx

We chose five cases that featured stem-immunity, immunity, and TLS-like structures on the same slide. In addition to one case from our 68 patient cohort (Patient #43), we identified four additional NSCLC cases in the MGH archives meeting these criteria (Patients #69, 70, 71, and 72). Slides were prepared using the semi-automated protocol as described in the manufacturer supplied protocol (Nanostring MAN-10151-01). 5um formalin-fixed paraffin-embedded tissue sections were baked at 65°C for 2 hours. We visualized *CXCL10* (Advanced Cell Diagnostics #311858-C2), *CCL19* (Advanced Cell Diagnostics #474368-C4), and *EPCAM* (Advanced Cell Diagnostics #310288-C3) expression using RNAscope probes to inform region of interest (ROI) selection. RNAscope probes amplified Cy3 (for visualization of *CXCL10*, Perkin Elmer #FP1046, 1:1500 dilution), Cy5 (for visualization of *CCL19*, Perkin Elmer #FP1171, 1:3000 dilution), and Alexa Fluor 594 (for visualization of *EPCAM,* ThermoFisher #B40957, 1:300 dilution) by tyramide amplification. A nuclear marker (Syto13, NanoString #121300303) was also visualized. Slides were profiled using the NanoString GeoMx Cancer Transcriptome Atlas (CTA, NanoString #121400101) and the Human TCR Alpha v1 panels in parallel.

Slides were loaded onto the GeoMx and scanned at 20x (following NanoString MAN-10152-01). ROI were manually selected as described in **Supplementary Fig. 5A**. In total, we collected 86 stem-immunity hub, 78 TLS, and 74 immunity hub ROIs across all patients. Library preparation was performed as described in the manufacturer supplied protocol (NanoString #MAN-10153-01), and sequencing was performed on the NextSeq500 platform.

GeoMx data were analyzed using the R packages NanoStringNCTools, GeomxTools, and GeoMxWorkflows (https://bioconductor.org). For CTA analyses, NanoString barcode counts were Q3 normalized (as described in manufacturer supplied protocol NanoString MAN-10154-01) for each ROI across all genes by patient. A mixed effect model was used to construct the volcano plots in **Fig. 4** and **Supplementary Fig. 5B-C** to account for slide-to-slide batch effects. CTA GSEA^60^ was performed using the R package fgsea (https://bioconductor.org) which calculates adjusted p-values and the Hallmark gene sets^32^.

### MERFISH

#### Data Generation

A MERSCOPE Gene Panel was designed using Vizgen’s Gene Panel Design Portal: https://portal.vizgen.com/. Briefly, a total of 484 genes were selected based on marker genes that commonly identify immune, stromal, and epithelial cells and expression programs in scRNAseq datasets (e.g. Reference^2^). 479 genes were assigned with a binary barcode for MERFISH imaging, while the remaining 5, due to their high expression level in cells, were imaged through sequential rounds of smFISH (**Supplementary Table 1O**).

We selected among PD-1-blockade responder cases based on whether they contained stem-immunity hubs and preference was given to blocks containing more tumor tissue to optimize for the number of cells per MERFISH run. Three of the cases were from the original cohort and the fourth was a newer case (#73) that became available after analysis of the original 68 patient cohort was complete. Formalin-Fixed Paraffin-Embedded (FFPE) human lung cancer samples were sectioned into 5um slices using a microtome and placed onto a 40mm diameter round MERSCOPE Slide. After drying at 55°C for 15 minutes, the tissue slices were deparaffinized and rehydrated by incubating in 100% ethanol, 90% ethanol and 70% ethanol, 5 minutes each. The samples were then de-crosslinked with heat-induced antigen retrieval at 90C for 15 minutes. Tissue sections were then blocked with Cell Boundary Blocking Buffer (Vizgen, PN 20300012) for 1 hour, stained with primary antibody mix (Vizgen, PN 20300010) diluted in Blocking Buffer at 1:100 for 1 hour, and then secondary antibody mix from Cell Boundary Staining Kit (Vizgen 20300011) for 1 hour. The sample was gel embedded, cleared, and hybridized with the MERSCOPE Gene Panel Mix at 37C for 36-48 hours by following Vizgen’s MERSCOPE FFPE Sample Preparation User Guide (https://vizgen.com/resources/merscope-formalin-fixed-paraffin-embedded-tissue-sample-preparation-user-guide/). Samples were washed twice with 5ml Formamide Wash Buffer at 47C for 30 minutes, before stained with DAPI and Poly T (Vizgen 20300021) for 15 minutes and washed with Formamide Wash Buffer at room temperature, away from light. Sample imaging was conducted on the MERSCOPE Platform (Vizgen 10000001).

#### Supervised Tissue ROI Selection

Output MERFISH images were reviewed using MERSCOPE Visualizer software and referenced against H&E sections. Low quality areas (e.g. necrosis, tissue detachment, and/or imaging artifact) were excluded from analysis. All tile and cell analyses described below were restricted to these high quality neoplastic regions.

#### Cell Type Segmentation

We performed cell segmentation using a combination of CellPose^61^ and Baysor^62^ algorithms. The CellPose algorithm was used to estimate cell shapes based on DAPI and a proprietary immunofluorescence cell surface marker cocktail (MERSCOPE User Guide 91600112 Rev B). Individual transcripts contained within the boundaries of CellPose segmented cells were assigned to their respective cells, while all others were assigned to background. We then used these initial cell assignments as the prior segmentation estimate for Baysor, an algorithm designed to identify cells in MERFISH data based on the spatial density of transcripts. We used the authors’ Julia implementation of Baysor (https://github.com/kharchenkolab/Baysor, Version 0.5.1) with the following parameters: min-molecules-per-segment = 2, min-pixels-per-cell = 15, scale-std = “100%”, iters=500, new-component-fraction = 0.3, prior-segmentation-confidence = 0.7, min-molecules-per-gene = 1, min-molecules-per-cell = 3, n-clusters = 1, new-component-weight = 0.2. For all other parameters, we used default values provided by the software. To run Baysor efficiently on our large datasets, we divided each image into tiles with at most 2,000,000 transcripts each, ran Baysor individually on each tile, and pooled the results from each tile into a global list of cells for each image.

#### Cell Type Labeling

We used a custom pipeline to analyze the individual cells segmented with the strategy above. We first performed quality control, removing all cells with fewer than 10 total transcripts. We then normalized and variance stabilized the read counts per cell by normalizing read counts to a fixed depth, adding 1, and applying the log transform. For the fixed depth, we used median total counts over all cells in the dataset, after QC filtering. We then performed a balanced PCA, as described in Korsunsky et al^63^, giving equal weight to each of the four datasets and to each region type (defined below). To account for batch effects, we used Harmony, with parameter theta=0.5, to correct for the effect of dataset identity on cells’ PCA embeddings. We then estimated a weighted nearest neighbor graph of the cells in harmonized PC space and used the graph to estimate a 2D UMAP embedding with the R uwot package^64^, parameters min_dist = 0.01 and spread = 0.22. We used the same graph to partition the cells by maximum modularity with the Leiden algorithm^59^, as implemented in the R igraph package^65^. We next identified cluster markers with differential expression analysis. To account for variable read depth and batch effects, we fit a Poisson generalized linear mixed effects model (GLMM) for each gene with the formula y_g ~ 1 + (1|cluster) + (1|cluster:batch) + (1|batch) + offset(logUMI). To do this, we used the presto-GLMM package (https://github.com/immunogenomics/presto/tree/glmm), which wraps GLMM estimation from lme4^66^ with random effect posterior variance estimation from arm^67^ to estimate the log fold-change (FC) and its variance of each gene in one cluster versus all other clusters. Finally, coarse-grained cell type labels were assigned to each cluster based on known lineage markers.

We repeated this full pipeline to subtype non-plasma lymphocytes and myeloid cells, which were difficult to disentangle in the initial coarse-grained clustering analysis of all cells. To protect against unwanted variation from contaminating transcripts, we removed genes that were significantly overexpressed (logFC>0, p<0.05) in epithelial cells, fibroblasts, mast cells, plasma cells, or vascular cells, and not overexpressed (logFC<0) in either lymphocyte or myeloid clusters in the coarse-grained marker analysis. We found that this strategy gave more emphasis to gene variation arising from intrinsic cell identity rather than confounding variation arising from neighboring cells or other forms of noise. Because we used a limited gene panel in this analysis, we performed an additional QC filtering step and removed cells within insufficient (<10 total transcripts) information. These cells were not given a fine-typed label and marked as poor-quality cells. On the remaining high quality cells, we performed normalization, PCA, Harmony integration, clustering and differential expression as described above. In order to ensure that we had clustered to a sufficient resolution, we checked that each top cluster marker was uniformly expressed throughout the cells in that cluster. Clusters whose top markers were expressed heterogeneously were further divided into subclusters using the same modularity clustering algorithm described above. Finally, a small number of clusters expressed marker genes associated with two cell types and could not be further subdivided to split cells into the two cell types. We labeled these cells as doublets and removed them from the final annotation.

#### Tissue segmentation and labeling

We performed unbiased tissue segmentation using a combination of grid-based binning and clustering analyses. We first divided each image into 2500um^2^ hexagon tiles and assigned transcripts to their respective overlapping tiles. To enforce smoothness in the gene expression profiles of neighboring tiles, we also added transcripts within 15um of a tile’s boundary to that tile. Note that with this strategy, one transcript may be assigned to up to 3 neighboring tiles. We then analyzed tiles using the clustering pipeline described in the Cell Type Labeling section above: remove low read tiles, normalize read counts with log(1+CP-median), PCA, Harmony, modularity clustering, and GLMM-based differential expression. We identified tile clusters based on differential expression of known markers: *CXCL9/10/11* and *CCL19* for stem-immunity, *CXCL9/10/11* (but not *CCL19*) for non-stem-immunity hubs, (**Supplementary Fig. 6F**), *PECAM1* and *VWF* for vascular hubs. All remaining tile clusters were labeled tumor. We next combined adjacent tiles with the same cluster labels into larger aggregates using deterministic graph partitioning based on weakly connected components using the igraph package^65^. We allowed for some uncertainty in tile label assignment by connecting similar tiles that were not directly adjacent but whose borders were within 50um of each other. We then pooled all transcripts from tiles inside an aggregate to define that aggregate’s gene expression profile. Final hub type markers were computed using GLMM-based differential expression on these aggregate gene expression profiles.

#### Colocalization analysis

Within each type of immunity hub, we analyzed the spatial organization of cells. Specifically, we asked whether two cells were physically adjacent to one another more often than expected by chance. To do this, we isolated all cells assigned to the same type of hub (e.g. stem-immunity) and identified their physically adjacent cells using Delaunay triangulation^68^. This strategy defined adjacent cells as cells whose centroids are within 30um of each other with no other cells located between their centroids. For each cell, which we will refer to as the index cell, we counted the number of neighbors of each type (e.g. 2 CD T cells, 1 B cell, 0 all others). We then tested, for each pair of cell types, whether one cell type (e.g. CD4 T cells) is more abundant in the neighborhood of another cell type (e.g. mregDCs) than expected by chance. To estimate the significance of the colocalization frequency of two cell types, we used spatial permutation, motivated by the approach of Keren et al^69^. Our strategy is as follows: fix the location of the index cell type, permute other cell type labels within spatial aggregates (as defined above in “Tissue Segmentation and Labeling”) 1000 times, compute the frequency of colocalization between index and other cell types under these random permutation, compute a variance for colocalization frequencies based on the null distribution from permutation colocalization frequencies, and finally use the variance to estimate a z-score for the observed colocalization frequency statistic. This permutation strategy is important for two reasons. First, it accounts for different densities of cell types, since more frequent cell types will have higher colocalization frequency, even under the spatial independence null model. Second, by permuting cell labels within aggregates only, we condition on the cell type composition of each aggregate. In practice, this approach resulted in better calibrated p-values. We derived statistical significance of colocalization frequencies using spatial permutation within spatial hubs, defined earlier as aggregates of adjacent tiles. This strategy did not work with our “tumor” region category, because the tumor regions were large and often spanned whole pieces of tissue. Spatial permutation works on spatially constrained regions and yields inflated p-values when applied to large regions. The intuition behind these inflated statistics is that spatial permutation is meant to test shuffling of labels in nearby cells. When we are allowed to shuffle labels of cells across the whole tissue (as in the tumor region), this assumption is broken and the null distribution is misspecified. We note that the calculation of colocalization is unlikely secondary to false positive T cell attribution because T cells have similar cell type signatures irrespective of whether or not they touch a *CXCL10*+ macrophage or mreg DC (**Supplementary Fig. 10**).

#### Cell with hub association testing

We tested the differential abundance of cell types within hub types to find which cell types are depleted and which are more abundant in which hub type (e.g. mregDCs in stem-immunity). To do this, we used each spatial aggregate, instead of individual tiles, as an independent observation, to ameliorate the confounding effect of spatial autocorrelation. Within each aggregate, we computed the average density of cell types as the average number of cells per tile. We then used linear mixed effects regression with lme4^66^ to compare the relative density of each cell type between two hub types. We fit a null model (density ~ 1 + (1|batch/hubType)) and an alternative model (density ~ 1 + hubType + (1|batch/hubType)) and computed the significance of the fixed effect with a likelihood ratio test between the alternative and null models. In these formulas, hubType is a two-level categorical variable that represents the two hub types we are testing (e.g. stem-immunity vs tumor). In the null model, we allow each library to have a different baseline density for a cell type and to have its own differential abundance between hub types. Thus, the likelihood ratio test will only be significant if the differential abundance is shared by multiple datasets and not only present in one or two.

## Acknowledgements

We thank the MGH Cancer Center Translational Imaging Core, MGH Pathology Department, members of the MGH spatial transcriptomics group, and members of the Hacohen lab.

This work was made possible by the generous support from Novartis as part of the MGH-Novartis Alliance. This work was also supported by public funds from NIH/NCI 1T32CA207021(J.H.C.) and R00CA259511 (K.P.), institutional funds from MGH Fund for Medical Discovery (J.H.C.) and MGH Krantz Stewardship (J.H.C.), and private funds from SU2C Phillip A. Sharp Award SU2C-AACR-PS-32 (K.P. and N.H.) and SITC/AstraZeneca Forward Fund (J.H.C.). N.H. is the David P. Ryan, MD Endowed Chair in Cancer Research, a gift from Arthur, Sandra, and Sarah Irving.

## Author contributions

J.H.C., L.T.N., M.S., M.M.-K., J.F.G., and N.H. conceptualized the study. J.H.C., L.T.N., V.J., J.D.P., K.Y., M.H., M.S.-F., and K.P. designed the RNA ISH/IF and IF only panels. J.H.C., M.S., V.J., and C.L. performed multiplex and GeoMx staining and imaging. J.H.C., L.T.N., M.S., V.J., and K.H.X. performed image analysis (including cell segmentation, cell phenotyping, and image co-registration). C.P. performed MERFISH imaging. J.H.C., L.T.N., M.S., P.M.R., S.F., S.S., A.M., M.P., I.Z., J.H., N.F.F., G.E., K.V.R., J.W.R., and M.A. performed computational analysis. J.H.C., C.S.N., C.B.M., J.L.B., M.Sakhi, G.M.B., M.M.-K., and J.F.G. performed clinical case finding and annotation. J.H.C., L.T.N., M.S., and N.H. wrote the manuscript. All authors edited the manuscript for intellectual content.

---

**Supplementary Figure 1.**
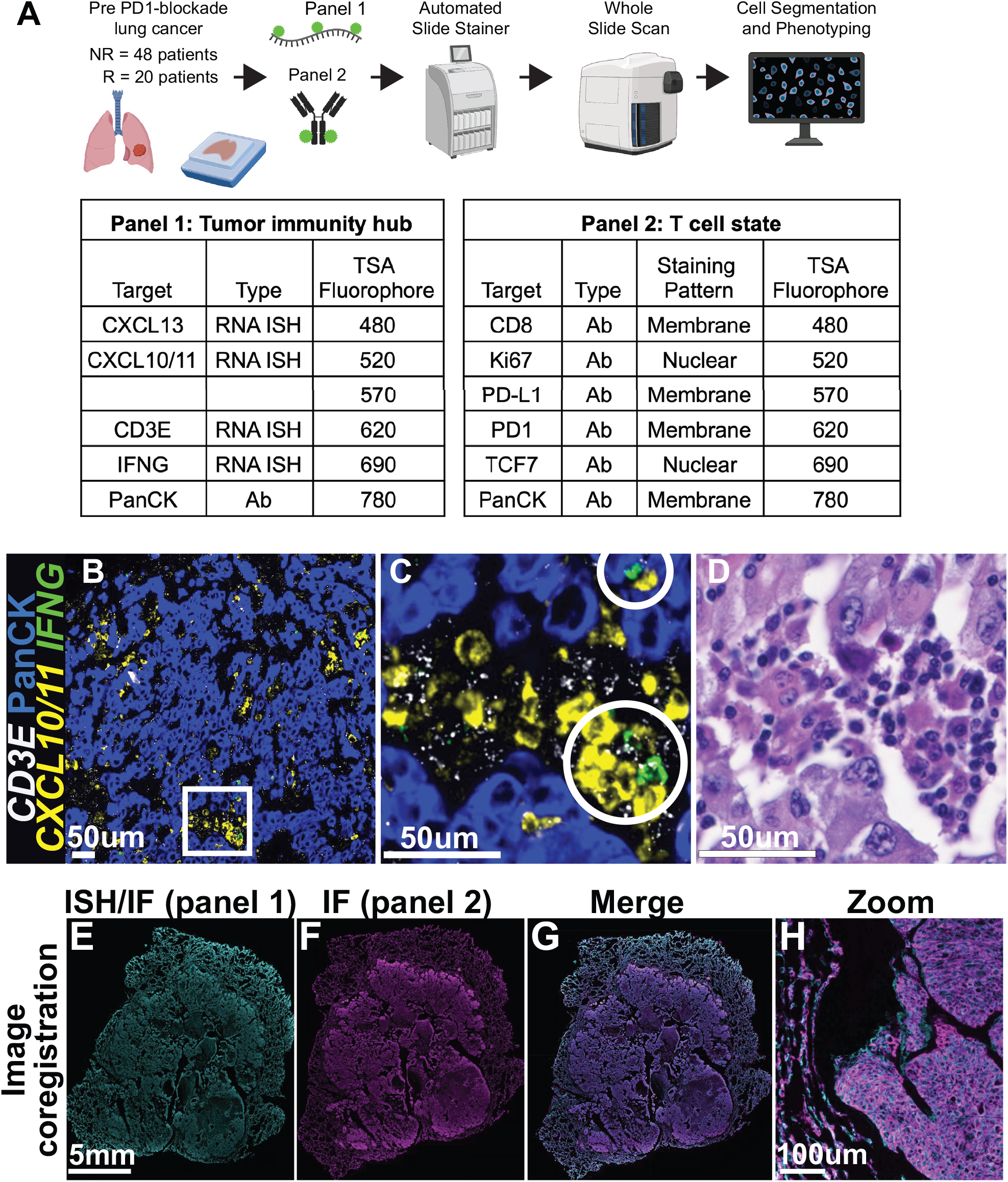
Workflow for investigating immunotherapy naive NSCLC patient samples. (A) Serial sections of pre-PD-1-blockade FFPE NSCLC samples were stained with two multiplex fluorescent panels for 68 patients (20 responders and 48 non-responders). Panel 1 uses RNA ISH/IF to identify the immunity hub components and panel 2 uses IF only to determine T cell states. Whole slide images were captured and analyzed by automated cell segmentation and phenotyping. (B) Representative low power image of a tumor (Patient #43) stained with the multiplexed RNA ISH/IF panel. (C) High power view from the boxed region in (B) showing an immunity hub. Cells positive for *IFNG* are marked by white circles. (D) High power view of H&E stained serial section from area matched to that in (C). (E-H) Paired RNA ISH/IF and IF only stained slides were coregistered for 46 of the patients. PanCK staining is shown in cyan for RNA ISH/IF and magenta for IF panel images.

**Supplementary Figure 2.**
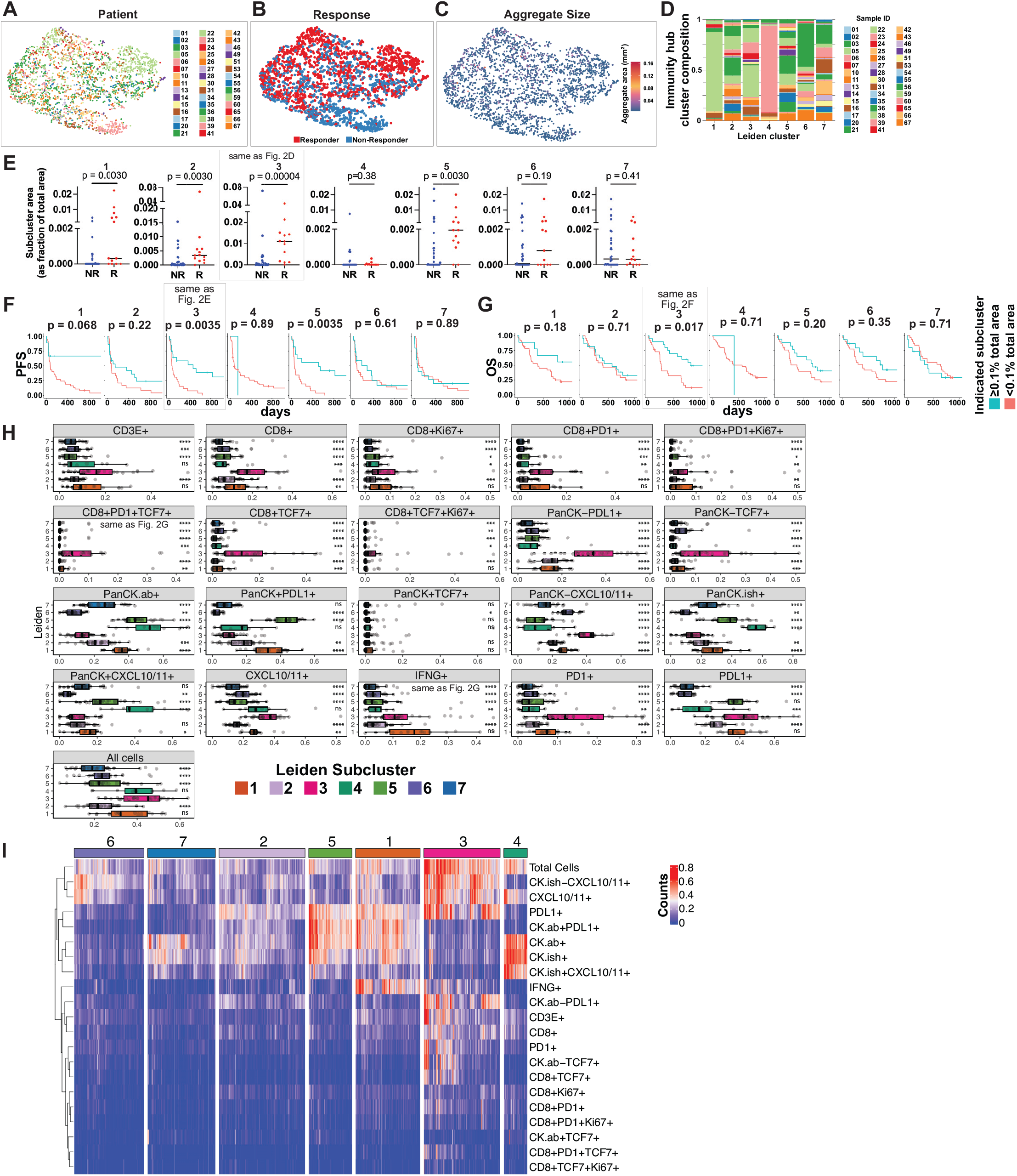
Leiden clustering of immunity hubs. (A-C) tSNE projection as in Fig. 2A colored by patient ID (A), patient response status (B), and immunity hub size (C). (D) Patient composition within each immunity hub Leiden subcluster. (E) Frequency of immunity hub subclusters expressed as a fraction of total tumor area for responders and non-responders. Mann-Whitney test p-value is shown. (F-G) Kaplan-Meier analysis of progression-free survival (PFS) and overall survival (OS) for each subcluster. Patients were classified as either above or below a threshold of 0.1% of total tumor area for each subcluster. Shown p-values are adjusted for multiple hypothesis testing using the Benjamini-Hochberg method (E-G). (H) Counts of indicated phenotypes within immunity hubs, as in Fig. 2G. Phenotype counts were normalized by immunity hub area and scaled from 0-1. Each point represents the mean value across all immunity hubs of a given Leiden subcluster for each patient having that subcluster. Statistical comparison was performed using an unpaired Mann-Whitney test relative to subcluster 3. Benjamini-Hochberg adjusted p-values shown: ns = not significant, *p<0.05, **p<0.01, ***p<0.001, and ****p<0.0001. (I) 0-1 scaled phenotype counts, as in (H), but for each of the 2712 hubs (columns).

**Supplementary Figure 3.**
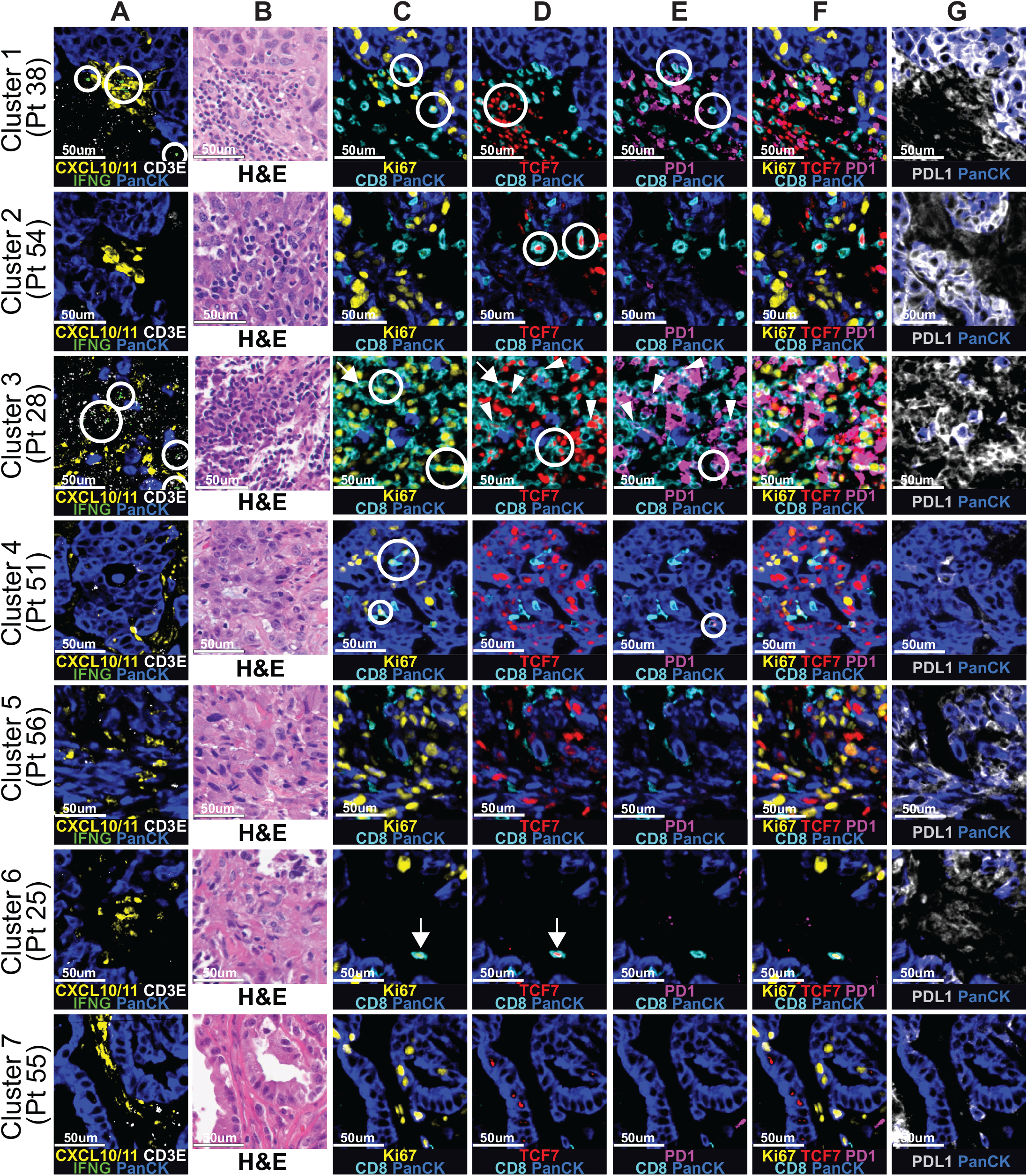
Representative images for each immunity hub Leiden subcluster. Each row features an immunity hub from a different patient. Aligned serial sections are shown for RNA ISH/IF panel (Column A), H&E (Column B), and IF only panel (Columns C-G). (A) *IFNG*+ cells (circles) are observed in immunity hubs. (B) H&E shows immunity hubs contain immune infiltrate abutting neoplastic cells. (C) Ki67+CD8+ (circles) and Ki67+TCF7+CD8+ cells (arrows) are found. (D) Ki67+TCF7+CD8+ (arrows), TCF7+CD8+ (circles), and TCF7+PD-1+CD8+ cells (arrowheads) are found. (E) PD-1+CD8+ (circles) and TCF7+PD-1+CD8+ cells (arrowheads) are found. (F) Composite image showing indicated channels. (G) PD-L1+ cells are found.

**Supplementary Figure 4.**
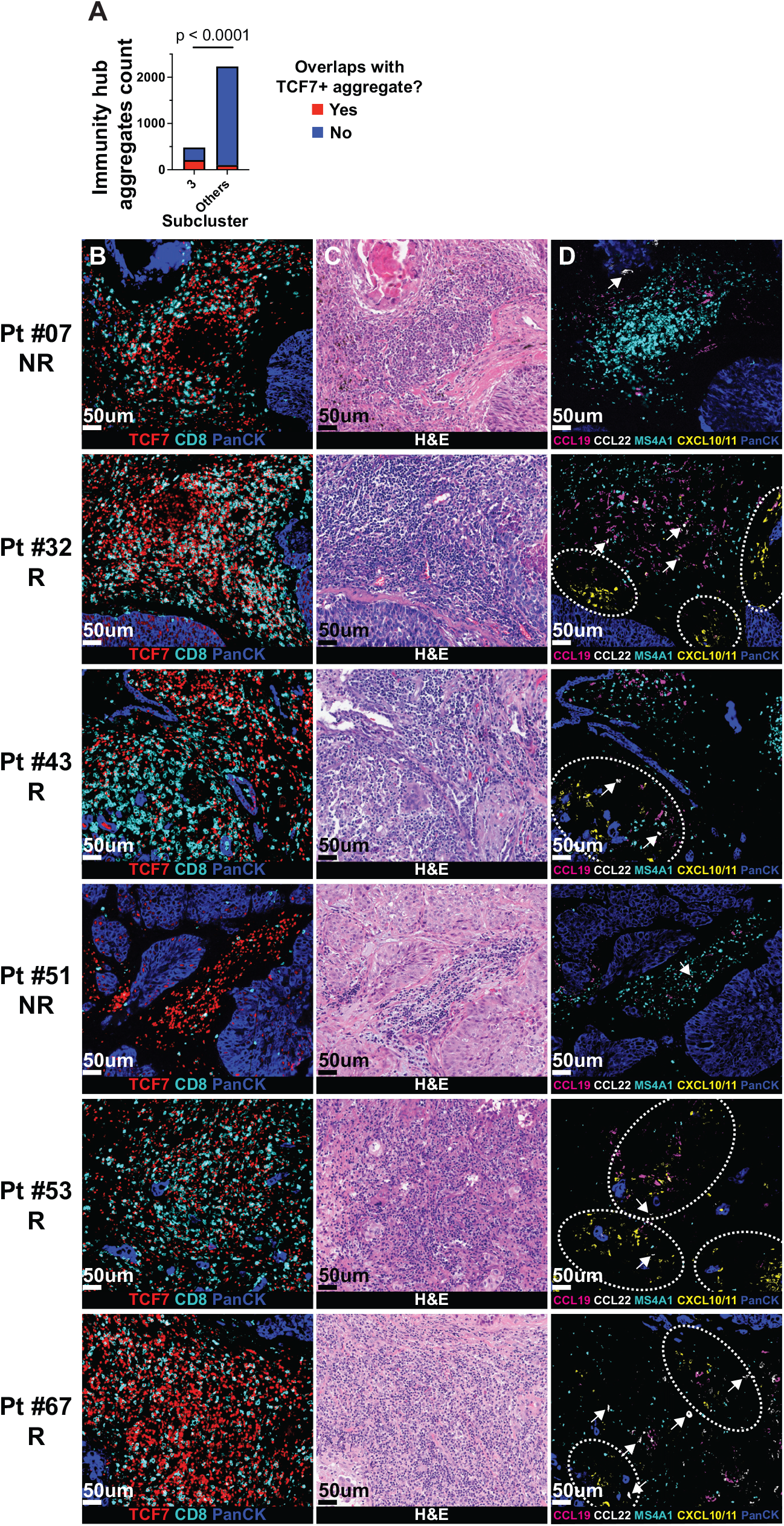
TCF7+ regions feature *CCL19*, *CCL22*, and variable *MS4A1* (encodes CD20), and frequently overlap with immunity hub subcluster 3. (A) Counts of contact (at least touching in a cardinal direction) between TCF7+ aggregates and subcluster 3 immunity hubs versus all other immunity hub subclusters (i.e. subclusters 1-2 and 4-7). Statistical comparison was performed using Fisher’s exact test. (B-D) Six cases (R=4, NR=2) from the 68 patient cohort with TCF7+ aggregates (ISH/IF panel staining shown in B) were stained by H&E (C) and an additional panel for *CCL19*, *CCL22*, *MS4A1*, *CXCL10/11*, and PanCK (D). Dashed ovals indicate areas with *CXCL10/11*+ cells. Arrows point to *CCL22*+ cells.

**Supplementary Figure 5.**
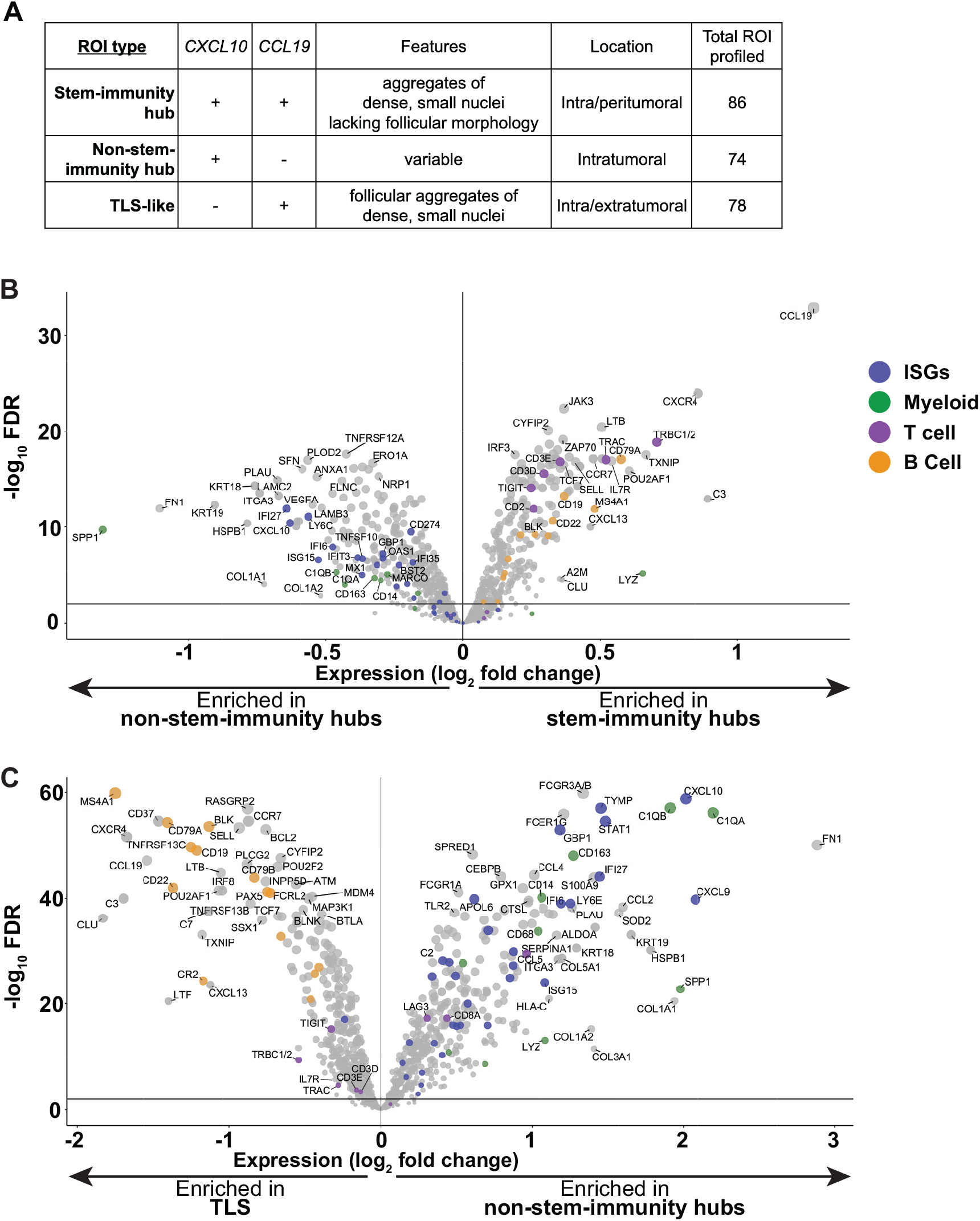
Transcriptional profile of stem-immunity hubs, non-stem-immunity hubs, and TLS using GeoMx. (A) Fluorescently labeled RNA ISH probes for *CXCL10* and *CCL19* on GeoMx slides guided region of interest (ROI) selection for transcriptional profiling. Histological features, region location, and number of regions profiled are noted. (B-C) Comparison of stem-immunity hub and non-stem-immunity hub (B) and non-stem-immunity hub and TLS (C) gene expression using GeoMx CTA assay. Volcano plots depict pooled ROIs across all 5 samples. Each dot represents one gene. Dot coloring is as follows: blue - ISGs, green - myeloid genes, purple - T cell genes, and orange - B cell genes.

**Supplementary Figure 6.**
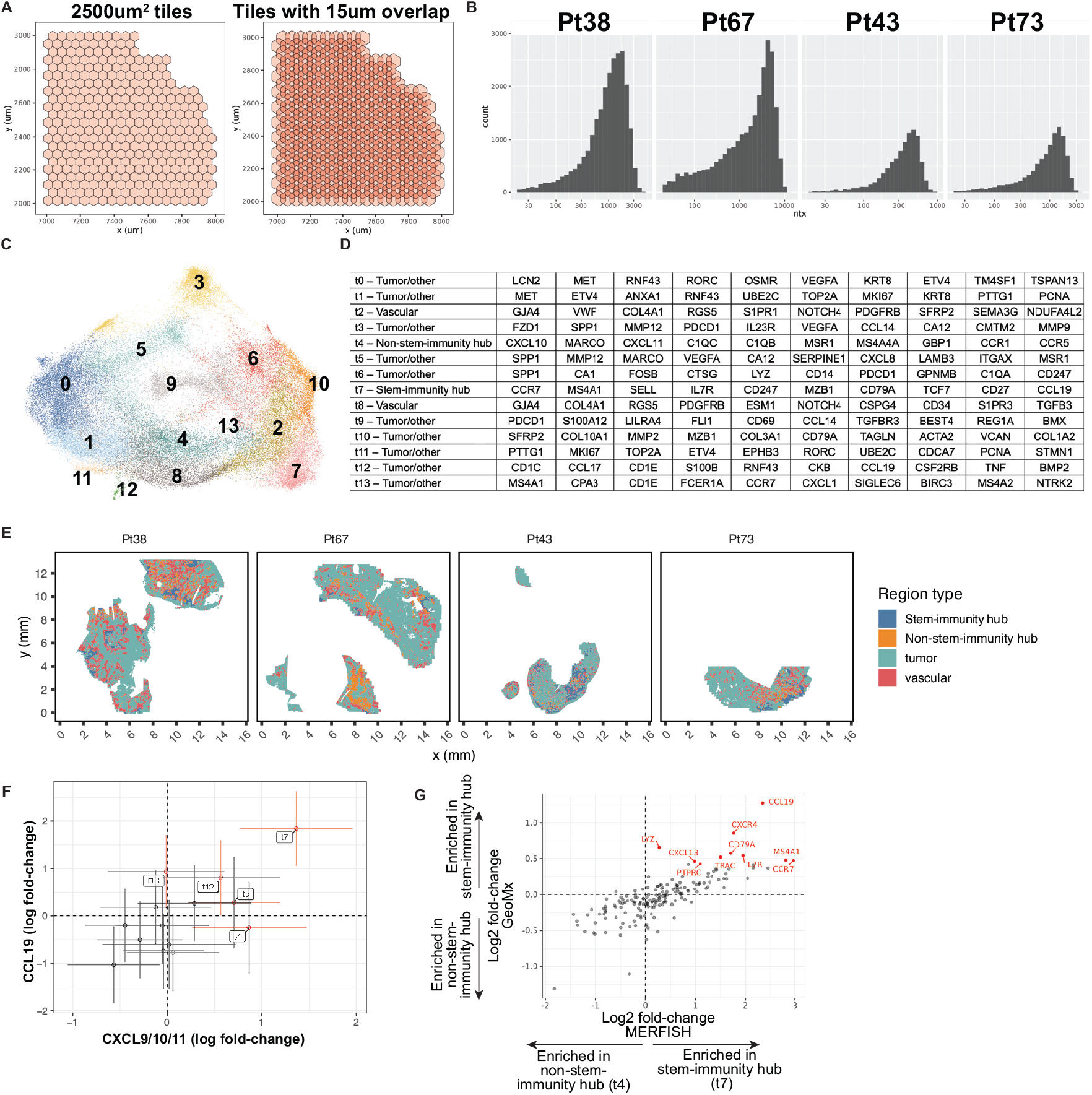
Tile-based analysis of MERFISH data reveals stem-immunity hubs in cluster t7. (A) The tile grid consisted of regular hexagons with area = 2500um^2^. To improve smoothness in clustering of tile tiles, hexagon tiles were dilated with a 15um buffer. (B) Transcripts per non-dilated 2500um^2^ tile by patient. (C) UMAP showing Louvain clustering of tiles. (D) Manual annotation of tile clusters using lists of top differentially-expressed genes for 14 cluster tiling solution. Cluster t7 marks stem-immunity hubs. (E) Spatial map of tumor for four MERFISH samples, with tiles colored by four region types. (F) Scatter plot of log fold-change for transcript abundance of *CCL19* and *CXCL9/10/11* for indicated tiles versus all other tiles. Tile cluster error bars colored red indicate significant enrichment (defined as FDR <20% and logFC>0) for *CXCL9/10/11* (horizontal) or *CCL19* (vertical). (G) Scatter plot of expression for 177 genes included in both GeoMx and MERFISH panels for manually annotated stem-immunity hubs (GeoMx) and tile cluster-defined stem-immunity hubs (t7, MERFISH) (r=0.78, p<1E-35). Axes indicate log2 fold-change of gene expression in stem-immunity versus non-stem-immunity hub. Each dot denotes one gene. Red dots denote the top 10 over-expressed genes in stem-immunity (versus non-stem-immunity hub) in the GeoMx dataset.

**Supplementary Figure 7.**
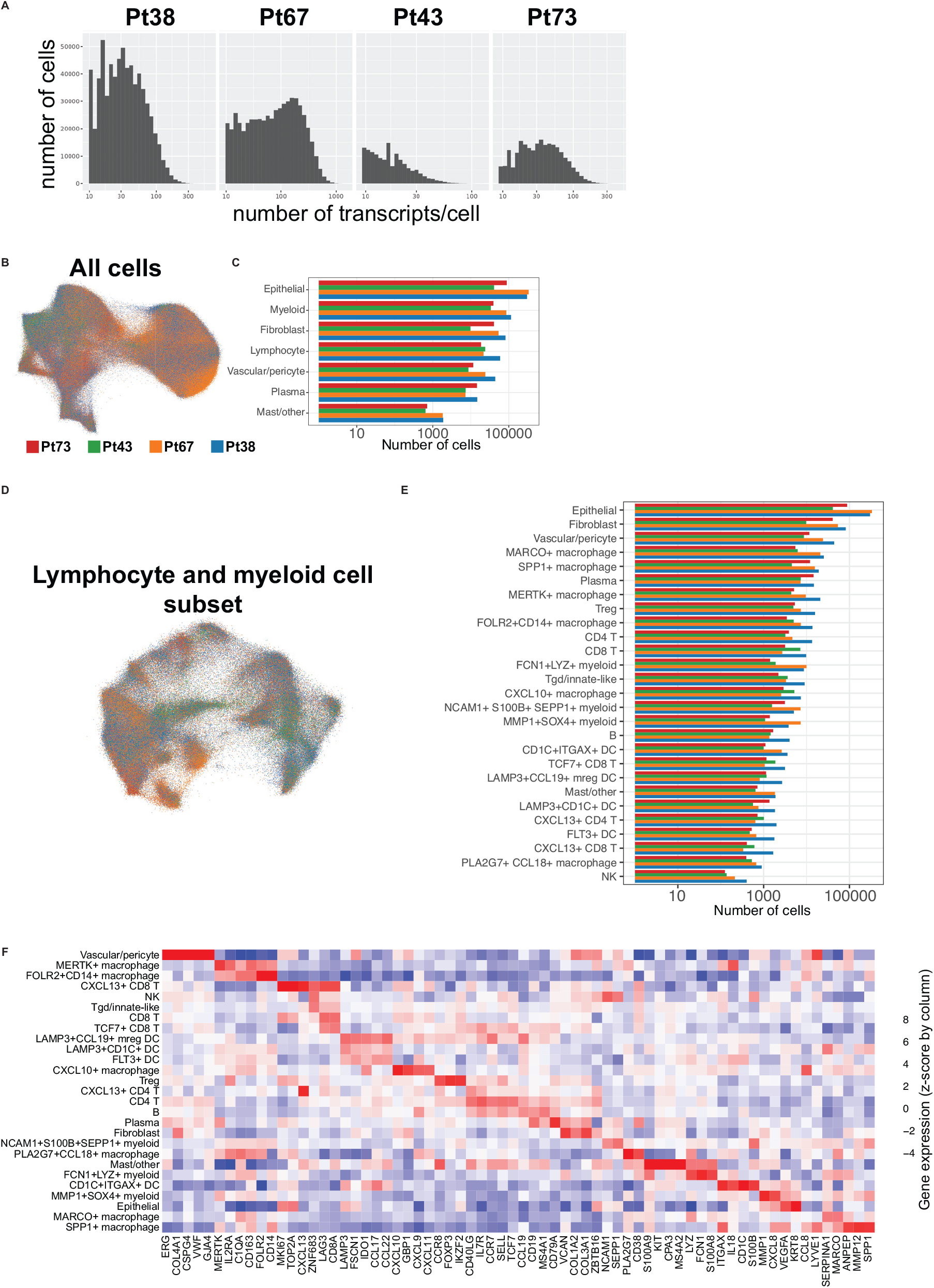
Cell-based clustering identifies many immune cell types in MERFISH data. (A) Histogram of transcripts per cell by patient. (B) UMAP of MERFISH data from Fig. 5C, colored by patient. (C) Cell counts by coarse cell type for each patient. (D) UMAP of immune cell MERFISH data from Fig. 5C, colored by patient. (E) Cell counts for fine immune clustering for each patient. (F) Heatmap including both coarse and fine cell clusters versus expression of key marker genes.

**Supplementary Figure 8.**
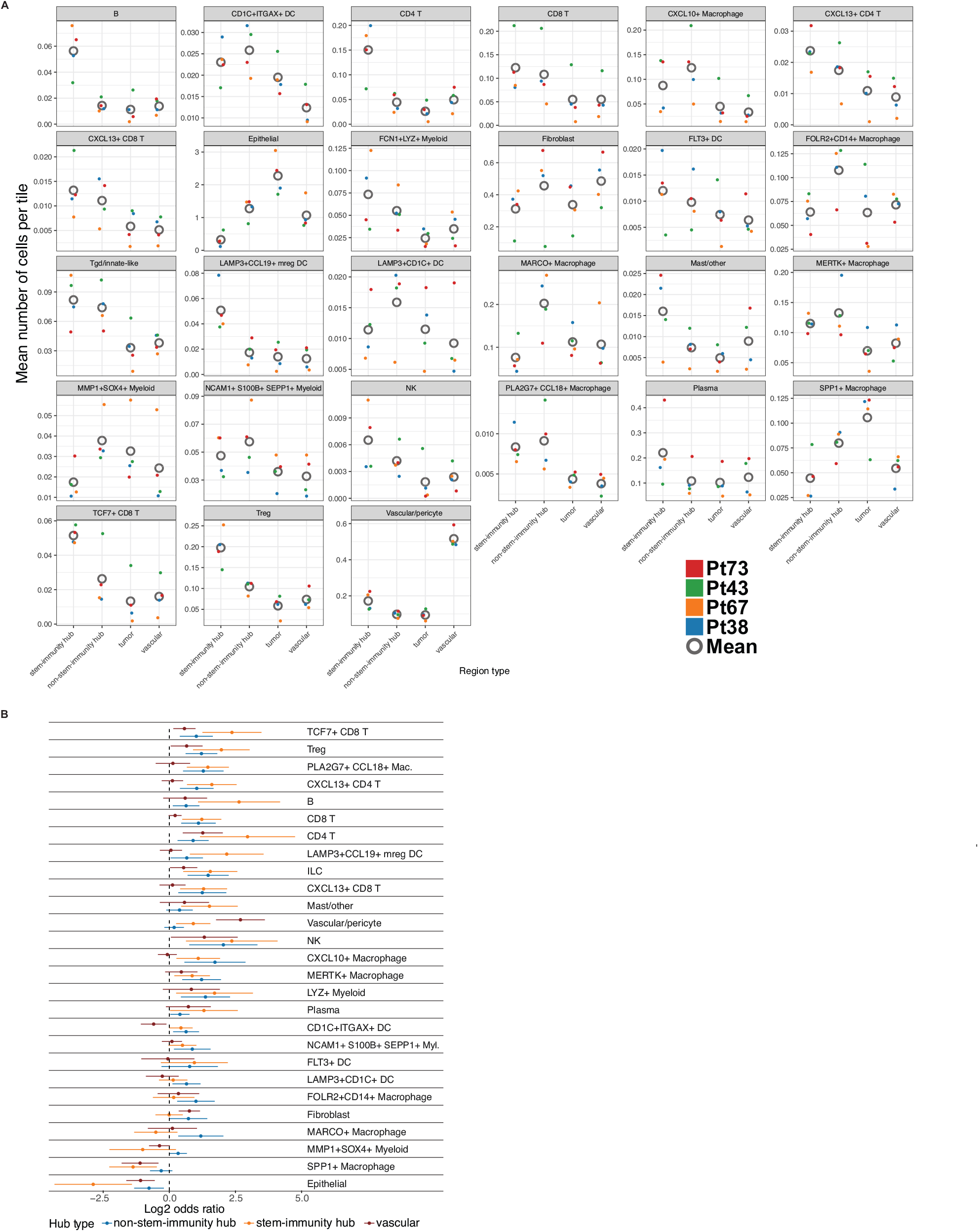
Cell-based analysis of MERFISH data reveals enrichment of mreg DCs, *TCF7*+ CD8 T cells, and other T cell populations within the stem-immunity hubs. (A) Mean number of cells per tile per aggregate for indicated clusters. Each dot represents one patient, colored by patient number. The open gray circle represents the mean cell per tile count for all aggregates of the indicated tile type across patients. (B) Forest plots depict the log2 odds ratio of the abundance (normalized to physical area) of a cell type in the indicated region type versus abundance in library-matched tumor areas. Error bars denote the 95% confidence interval.

**Supplementary Figure 9.**
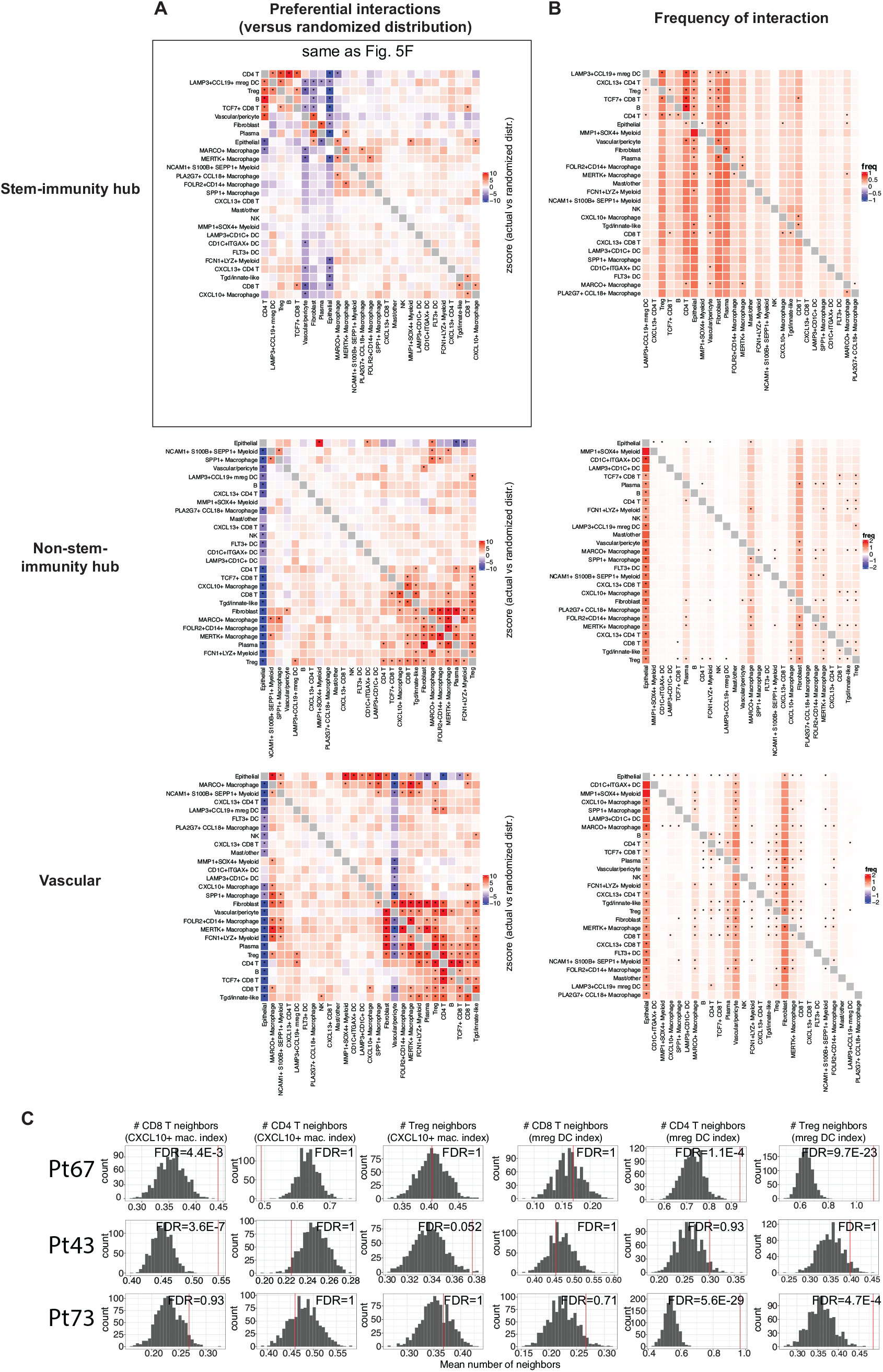
Non-random cell-cell interactions across tile neighborhood types. (A) Neighboring cell analysis for cells within indicated tile region types. For each plot, indicated cluster position (row) is held constant and enrichment with partner (column) is quantified versus scrambled distribution. Mean z-scores across all four samples are shown. Two-tailed p-values were calculated from the mean z-scores and FDRs computed with the Benjamini-Hochberg procedure across all p-values. FDR<0.05 are denoted with an asterisk. (B) Absolute frequency of interactions (mean of the mean frequencies across the four samples) for a given index cell (row) and partner cell type (column). (C) As in Fig. 6B, the red line on histograms shows how frequently specific T cell types are observed to neighbor the indicated myeloid cell type within the stem-immunity hubs for indicated patients. Gray bars denote random distribution (expected distribution based on cell type frequency).

**Supplementary Figure 10.**
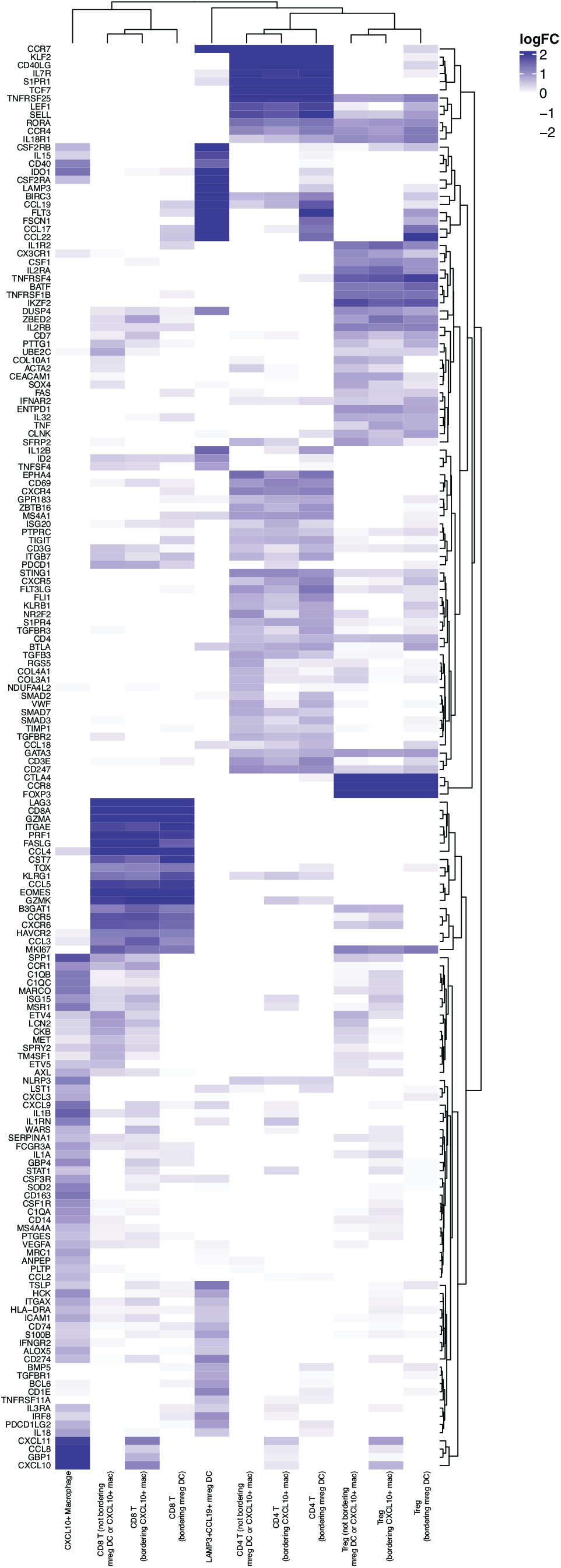
Gene signatures of T cells neighboring mreg DCs and *CXCL10*+ macrophages. Heatmap of gene expression of CD4 T cells, CD8 T cells, and Tregs, subdivided into three groups reflecting their neighborhood composition: adjacent to at least one *LAMP3*+*CCL19*+ mreg DC, adjacent to at least one *CXCL10*+ macrophage, or adjacent to neither. T cells adjacent to both were rare and thus not included in this analysis. Only genes significantly differentially overexpressed (adj. p<0.05 & logFC>0.5) in one of the rows is shown.

